# DNAH3 deficiency causes flagellar inner dynein arm loss and male infertility in humans and mice

**DOI:** 10.1101/2024.02.19.580977

**Authors:** Xiang Wang, Gan Shen, Yihong Yang, Chuan Jiang, Tiechao Ruan, Xue Yang, Liangchai Zhuo, Yingteng Zhang, Yangdi Ou, Xinya Zhao, Shunhua Long, Xiangrong Tang, Tingting Lin, Ying Shen

## Abstract

Axonemal protein complexes, including the outer and inner dynein arms (ODA/IDA), are highly ordered structures of the sperm flagella that drive sperm motility. Deficiencies in several axonemal proteins have been associated with male infertility, which is characterized by asthenozoospermia or asthenoteratozoospermia. Dynein axonemal heavy chain 3 (DNAH3) resides in the IDA and is highly expressed in the testis. However, the relationship between DNAH3 and male infertility is still unclear. Herein, we identified biallelic variants of *DNAH3* in four unrelated Han Chinese infertile men with asthenoteratozoospermia through whole-exome sequencing (WES). These variants contributed to deficient DNAH3 expression in the patients’ sperm flagella. Importantly, the patients represented the anomalous sperm flagellar morphology, and the flagellar ultrastructure was severely disrupted. Intriguingly, *Dnah3* knockout (KO) male mice were also infertile, especially showing the severe reduction in sperm movement with the abnormal IDA and mitochondrion structure. Mechanically, nonfunctional DNAH3 expression resulted in decreased expression of IDA-associated proteins in the spermatozoa flagella of patients and KO mice, including DNAH1, DNAH6, and DNALI1, the deletion of which has been involved in disruption of sperm motility. Moreover, the infertility of patients with *DNAH3* variants and *Dnah3* KO mice could be rescued by intracytoplasmic sperm injection (ICSI) treatment. Our findings indicated that *DNAH3* is a novel pathogenic gene for asthenoteratozoospermia and may further contribute to the diagnosis, genetic counseling, and prognosis of male infertility.

## Introduction

Infertility is a global public health and social problem that affects approximately one in six couples worldwide (1). Male infertility, which accounts for half of infertile cases, is a multifactorial disease with common phenotypes, including oligo/azoospermia (poor sperm count or absence of spermatozoa); teratozoospermia (aberrant sperm morphology); asthenozoospermia (weakened sperm motility); and a combination of these phenotypes, such as asthenoteratozoospermia, oligoasthenozoospermia, oligoteratozoospermia and oligoasthenoteratozoospermia (2, 3).

Asthenoteratozoospermia is one of the most common phenotypes of male infertility, and genetic factors have been established as the predominant cause of asthenoteratozoospermia. Multiple morphological abnormalities of the flagella (MMAF), a subtype of asthenoteratozoospermia, characterized by a mosaic of abnormalities of the flagellar morphology, including absent, short, coiled, bent and/or irregular flagella, is almost always caused by genetic defects (4, 5). To date, more than 40 genes have been identified as pathogenic genes of MMAF, but these genes can only explain approximately 60% of MMAF-affected cases (6–9). Therefore, the genetic basis of the remaining cases is still unknown.

The motility of a sperm is driven by its rhythmically beating flagella, and at the center of the flagella lies a conserved axonemal structure, containing the “9 + 2” microtubular arrangement: a ring of nine microtubule doublets (MTDs) surrounding a central pair (CP) of singlet microtubules. Each MTD consists of an A tubule and a B tubule, and the outer (ODA) and inner (IDA) dynein arms are anchored along the A tubule (10). The ODA and IDA are ATPase-based protein complexes that drive the movement between the A tubule and the neighboring B tubule of the next doublet, producing the original force for sperm motility (11, 12). Structural and functional abnormalities of the ODA and IDA have been demonstrated to cause male infertility associated with asthenozoospermia and/or asthenoteratozoospermia (4, 13, 14).

The dynein axonemal heavy chain (DNAH) family comprises a series of proteins (DNAH1–3, DNAH5–12, and DNAH17) that are precisely assembled with other axonemal dynein motor proteins in the ODAs and IDAs of sperm flagella and motile cilia (15–17). In humans, DNAH1, DNAH2, DNAH6, DNAH7, DNAH8, DNAH10, DNAH12 and DNAH17 are highly expressed in the testis, and deficiency of these proteins has been demonstrated to cause MMAF-associated asthenoteratozoospermia (18–25). DNAH3 is an evolutionarily conserved IDA-associated protein and is highly expressed in testes of humans and mice (26). Deficient DNAH3 has been shown to impair sperm motility in *Drosophila* and cattle (27, 28). In humans, *DNAH3* has been identified as a novel breast cancer candidate gene (29). However, the role of DNAH3 in male reproduction in humans and mice remains largely unknown.

In the present study, we identified four biallelic variations in *DNAH3* in four unrelated Han Chinese patients with asthenoteratozoospermia using whole-exome sequencing (WES). The spermatozoa of the patients showed extremely reduced sperm motility and a high proportion of sperm tail defects characterized by the MMAF phenotype. We further generated *Dnah3* knockout (KO) mice, and the male KO mice expectedly showed aberrations in sperm movement, flagellar IDA, and mitochondrion. Moreover, the absence of DNAH3 led to decreased expression of other IDA-associated proteins, including DNAH1, DNAH6 and DNALI1. Importantly, good outcomes of intracytoplasmic sperm injection (ICSI) treatment were observed in *DNAH3*-deficient patients and *Dnah3* KO mice. This study revealed *DNAH3* as a novel pathogenic gene of asthenoteratozoospermia, and the findings provide valuable suggestions for the clinical diagnosis and treatment of male infertility.

## Results

### Identification of biallelic pathogenic variants of *DNAH3* in four unrelated infertile men

In the present study, we employed whole-exome sequencing (WES) to identify potential candidate variants associated with primary asthenoteratozoospermia. After comprehensive filtering and screening, we identified 98, 101, 67 and 91 candidate variants for Patient 1, Patient 2, Patient 3 and Patient 4, respectively (**Table S1**). To refine these candidate variants, we excluded those whose corresponding genes were not expressed in the human or mouse testis, were associated with diseases unrelated to male infertility, or were monoallelic variants. Ultimately, only bi-allelic variants in *DNAH3* (NG_052617.1, NM_017539.2, NP_060009.1) remained, suggesting as the pathogenic variants responsible for the infertility of the patients : a compound heterozygous mutation of c.3590C>T (p.Pro1197Leu) and c.3590C>G (p.Pro1197Arg) in Patient 1, a homozygous missense mutation of c.4837G>T (p.Ala1613Ser) in Patient 2, a compound heterozygous mutation of c.5587del (p.Leu1863*) and c.10355C>T (p.Ser3452Leu) in Patient 3 and a compound heterozygous mutation of c.2314C>T (p.Arg772Trp) and c.4045G>A (p.Asp1349Asn) in Patient 4 (**Figure 1A**). Importantly, routine semen analysis revealed that all patients showed extremely reduced sperm motility and a high proportion of sperm tail defects (**Table 1**). These variants either were not recorded or had an extremely low frequency in East Asian population in multiple public population databases, including the ExAC browser, GnomAD and the 1000 Genomes Project, and were predicted to be potentially deleterious by SIFT (https://sift.bii.a-star.edu.sg/), PolyPhen-2 (http://genetics.bwh.harvard.edu/pph2/), MutationTaster (https://www.mutationtaster.org/), and CADD (https://cadd.gs.washington.edu/) (**Table 2**) (30–33). Next, Sanger sequencing confirmed these variants in the probands, and their fertile parents carried the heterozygous variants (**Figure 1A**). Moreover, the variant sites are localized in several domains of the DNAH3 protein and are highly conserved across species (**Figure 1B**).

**Figure 1.**
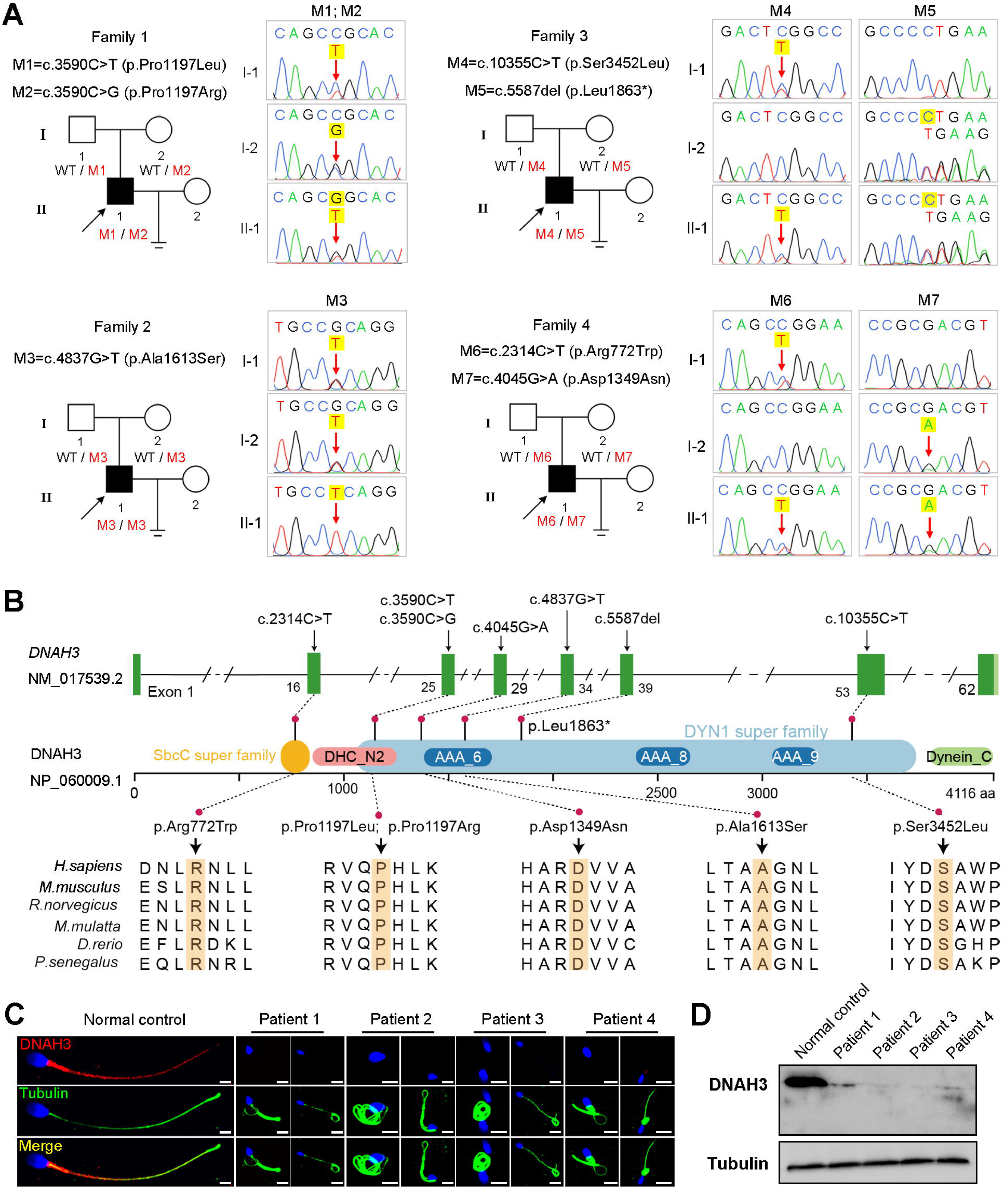
Identification of biallelic pathogenic variants in *DNAH3* from four unrelated infertile families. (**A**) Pedigrees of four families affected by *DNAH3* variants (M1–M7). Black arrows indicate the probands in these families. (**B**) Location of the variants and conservation of affected amino acids in DNAH3. Black arrows indicate the position of the variants. (**C**) Immunofluorescence staining of DNAH3 in sperm from the patients and normal control. Red, DNAH3; green, α-Tubulin; blue, DAPI; scale bars, 5 μm. (**D**) Western blotting analysis of DNAH3 expressed in spermatozoa from the patients and normal control.

**Table 1.** Semen analysis of the patients in the present study.

|  |  | Patient 1 | Patient 2 | Patient 3 | Patient 4 | Reference |
| --- | --- | --- | --- | --- | --- | --- |
| **Semen parameters** | Semen volume (ml) | 4.3 | 3.3 | 0.8 | 4.6 | ≥1.5 |
|  | Semen concentration (10<sup>6</sup>/ml) | 2.0 | 0.5 | 11.0 | 21.0 | ≥15.0 |
|  | Motility (%) | 6.0 | 3.0 | 0 | 0 | ≥40.0 |
|  | Progressive motility (%) | 0 | 2.3 | 0 | 0 | ≥32.0 |
| **Sperm morphology** | Normal (%) | 1.3 | 1.1 | 1.8 | 1.2 | ≥4.0 |
|  | Tail defects (%) | 91.3 | 87.5 | 96.5 | 97.5 | - |

-, not applicable.

**Table 2.**
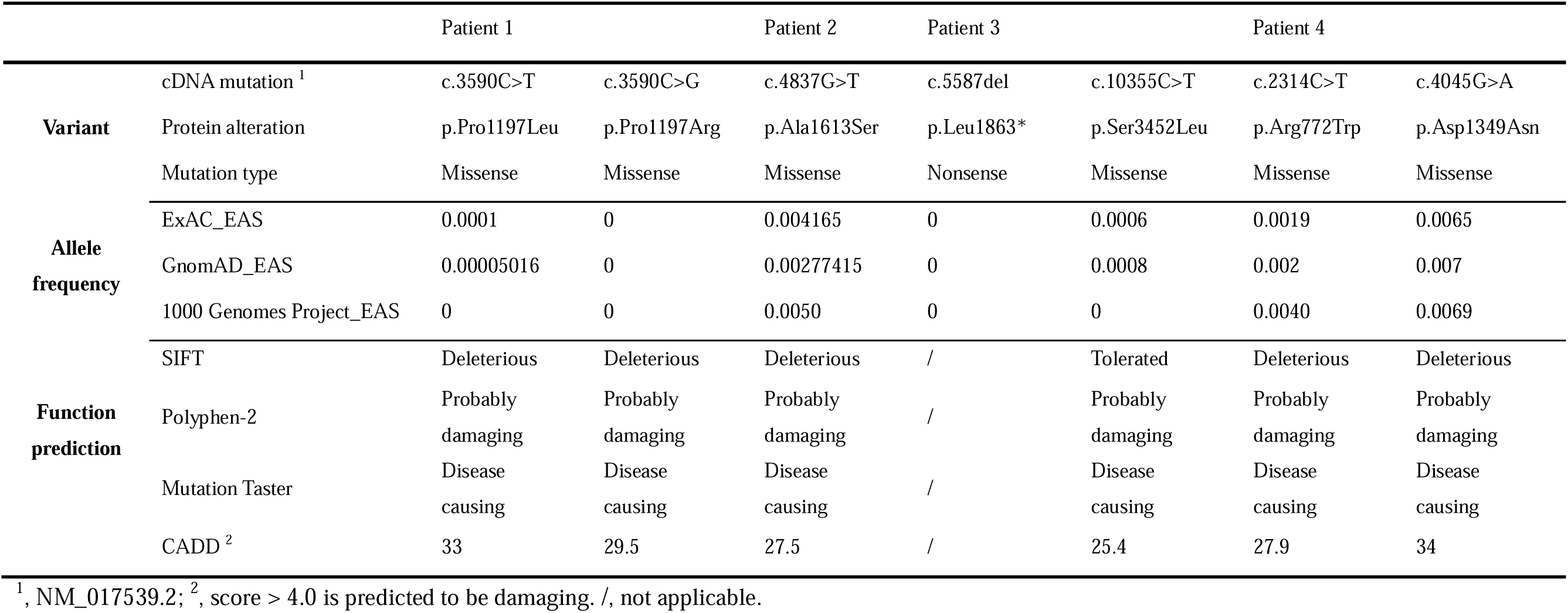
Variants analysis of the patients in the present study.

Strikingly, immunofluorescence staining revealed that DNAH3 was exclusively resided in the tail and concentrated in the midpiece of control sperm. However, the fluorescence signal of DNAH3 was hardly detected in the patients’ spermatozoa (**Figure 1C**). Additionally, subsequent western blotting analysis yielded consistent results with immunofluorescence staining, indicating that these variants led to disrupted expression of DNAH3 (**Figure 1D**). These results suggested that biallelic variants in *DNAH3* disrupted DNAH3 expression and might be responsible for the infertility of the four patients.

### Asthenoteratozoospermia phenotype is observed in patients with *DNAH3* variants

We next investigated the aberrant sperm morphology of the patients using Papanicolaou staining and SEM analysis. Notably, the tails of sperm from the patients exhibited a typical phenotype associated with MMAF, including coiled, short, bent, irregular, and/or absent flagella (**Figure 2A and B**, **Figure S1A**). In addition, a fraction of defects in the sperm head were also present in the patients’ sperm (**Figure 2B**).

**Figure 2.**
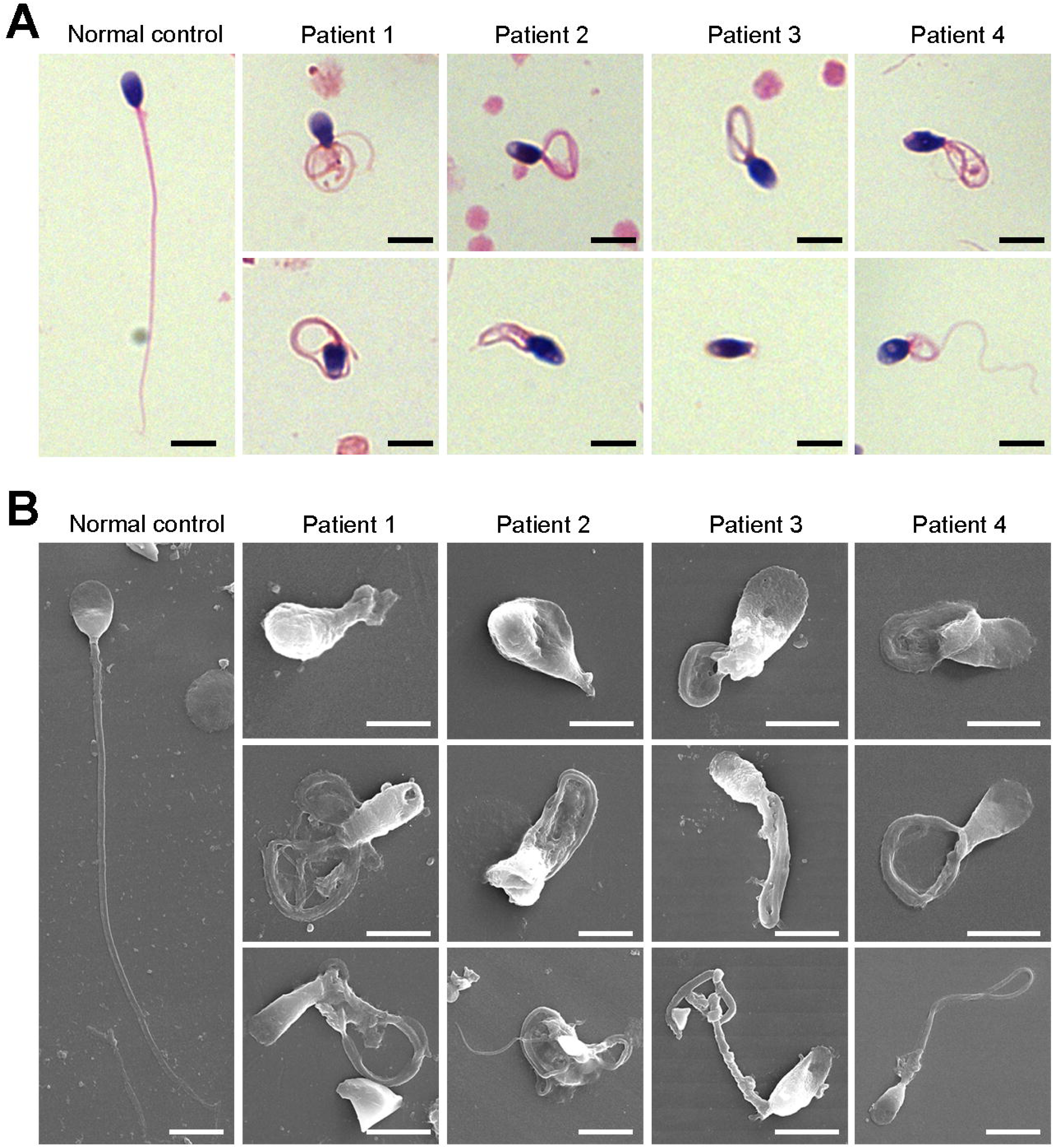
Defects in sperm morphology of the patients harboring *DNAH3* variants. (**A**, **B**) Abnormal sperm morphology was observed through Papanicolaou staining (A), and SEM analysis (B) compared to normal control. Scale bars, 5 μm.

TEM was employed to determine the ultrastructure of the sperm from the patients. Compared to the integrated and well-organized “9L+L2” axonemal arrangement of the sperm flagella from the normal control, spermatozoa from the patients showed absent or disordered CPs, MTDs, and outer dense fibers (ODFs) in different regions of the flagella (**Figure 3A**, **Figure S1B**). Interestingly, the IDAs of sperm flagella of the patients were hardly captured compared to the control (**Figure 3A**). Additionally, in the midpiece of sperm flagella of the patients, dissolved mitochondrial material was also observed evidently under TEM (**Figure 3A**). We next conducted immunofluorescence staining to label the mitochondria of patients’ sperm with TOM20, a subunit of the mitochondrial import receptor. Remarkably, in contrast to the robust TOM20 signals observed in the normal control, the TOM20 signals in the sperm from the patients were considerably diminished, indicating a disrupted mitochondrial function (**Figure 3B**). Together, these data suggested that *DNAH3* may function in sperm flagellar development, and loss-of-function variants were associated with MMAF in humans.

**Figure 3.**
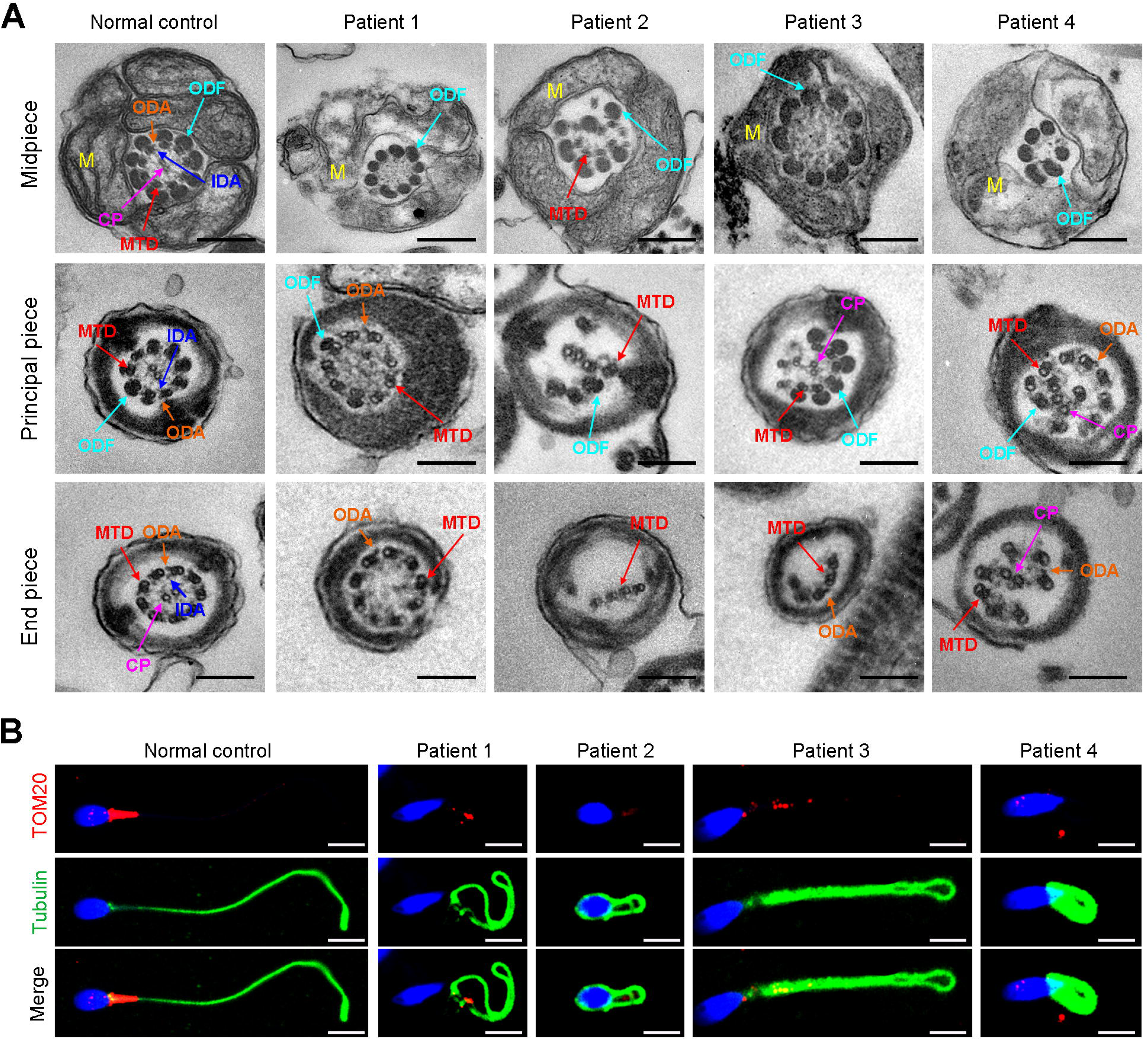
Ultrastructural and mitochondrial defects in sperm from infertile men with *DNAH3* variants. (**A**) TEM analysis of sperm obtained from a normal control and patients harboring *DNAH3* variants. Cross-sections of the midpiece, principal piece and endpiece of sperm from normal control showed the typical ‘‘9 + 2’’ microtubule structure, and an IDA and an ODA were displayed on the A-tube of each microtubule doublet. Cross-sections of the midpiece, principal piece and endpiece of sperm from the patients displayed absent or disordered CPs, MTDs and ODFs, as well as an evident missing of the IDAs in different pieces of the flagella. M, mitochondria sheath; ODF, outer dense fiber; MTD, microtubule doublets; CP, central pair; IDA, inner dynein arms; ODA, outer dynein arms. Scale bars, 200 nm. (**B**) Immunofluorescence staining of TOM20 in sperm from the patients and normal control. Red, TOM20; green, α-Tubulin; blue, DAPI; scale bars, 5 μm.

### DNAH3 is exclusively expressed in the sperm flagella of humans and mice

To further understand the function of DNAH3 in male reproduction, we explored the expression pattern of DNAH3 in humans and mice. qPCR results revealed that *Dnah3* was predominantly expressed in the mouse testis (**Figure S2A**). Moreover, when observing the expression of *Dnah3* in testes from mice at different postnatal days, we found that *Dnah3* expression was significantly elevated beginning on postnatal Day 22, peaked at postnatal Day 30, and maintained a stable expression level thereafter (**Figure S2B**). In addition, germ cells at different stages were isolated from the testes of humans and mice and were stained with anti-DNAH3 antibody. The results showed that DNAH3 was expressed in the cytoplasm of spermatocytes and spermatogonia and then obviously in the flagellum of early and late spermatids (**Figure S3A and B**). These expression data suggest that DNAH3 may play an important role in sperm flagellar development during spermatogenesis in humans and mice.

### Deletion of *Dnah3* causes male infertility in mice

Considering the absent expression of DNAH3 in the patient sperm, we generated *Dnah3* KO mice using CRISPRLCas9 technology to further confirm the essential role of DNAH3 in spermatogenesis (**Figure S4A**). PCR, qPCR, and immunofluorescence staining were used to confirm that *Dnah3* was null in KO mice (**Figure S4B-E**). The *Dnah3* KO mice survived without any evident abnormalities in development and behavior. H&E staining further revealed that there were no histological differences in the lung, brain, eye, or oviduct between wild-type (WT) and *Dnah3* KO mice (**Figure S5A**). In addition, no obvious abnormalities in ciliary development were observed in these organs in KO mice compared to WT mice (**Figure S5B**). The *Dnah3* KO female mice were fertile with normal oocyte development (**Figure S6A**). However, the *Dnah3* KO male mice were completely infertile (**Figure 4A**). We next examined the testis and epididymis of *Dnah3* KO male mice to elucidate the etiology of infertility. There was no detectable difference in the testis/body weight ratio of *Dnah3* KO mice when compared to WT mice (**Figure S6B**).

**Figure 4.**
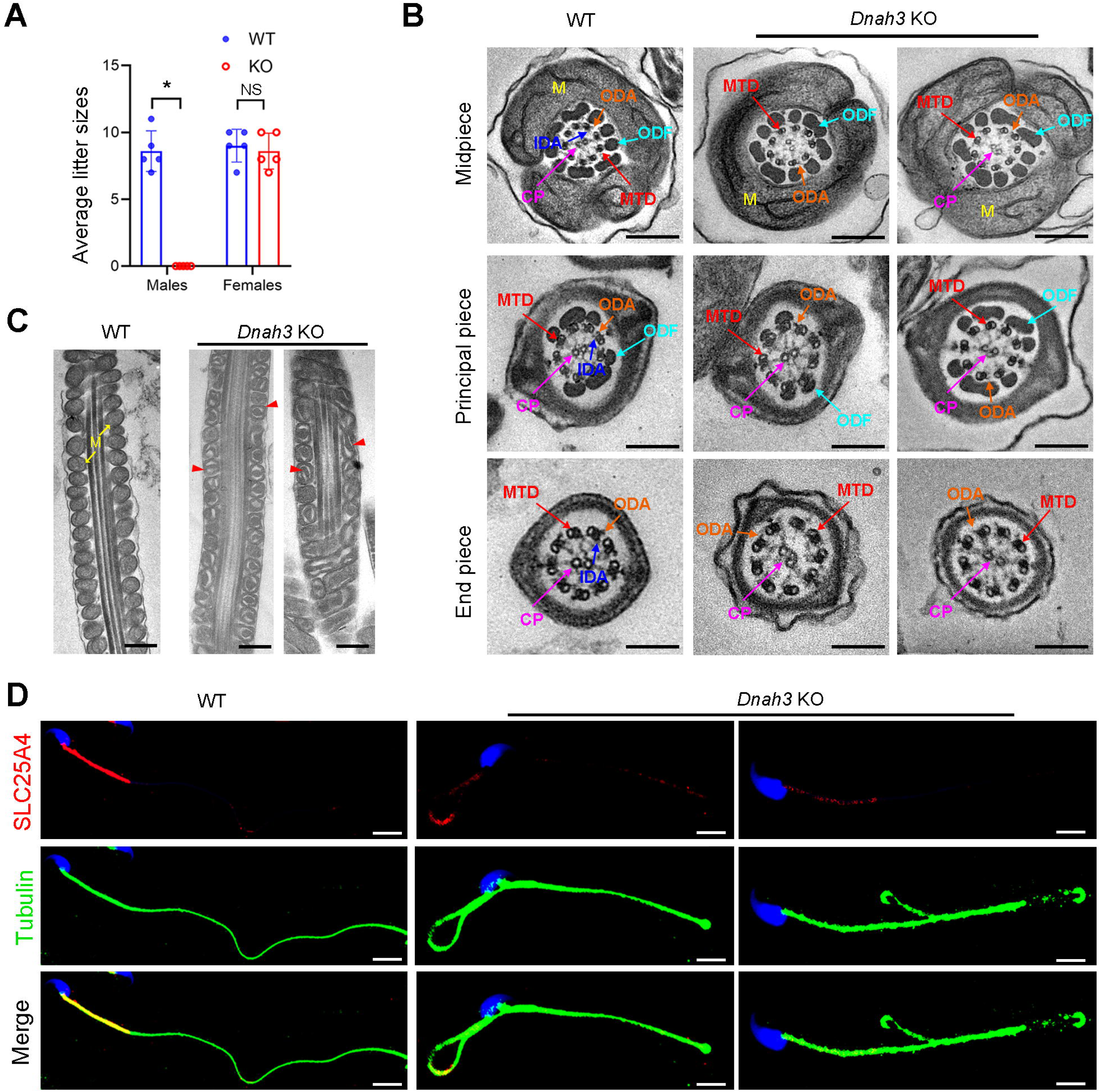
*Dnah3* KO male mice are infertile. (**A**) Fertility of *Dnah3* KO mice. The KO male mice were infertile (n = five biologically independent WT mice or KO mice; Student’s *t* test; *, P < 0.05; NS, no significance; error bars, s.e.m.). (**B**) TEM analysis of the cross-sections of spermatozoa from *Dnah3* KO mice revealed an obvious absence of IDAs in different pieces of the flagella compared to WT mice. M, mitochondrion sheath; ODF, outer dense fiber; MTD, microtubule doublet; CP, central pair; IDA, inner dynein arm; ODA, outer dynein arm. Scale bars, 200 nm. (**C**) Disrupted mitochondria were observed in spermatozoa tail from *Dnah3* KO mice by TEM analysis. The yellow arrows indicate the normal mitochondria. The red arrowheads indicate the dilated intermembrane spaces and dissolved mitochondrial material. M, mitochondrion sheath. Scale bars, 200 nm. (**D**) Immunofluorescence staining of SLC25A4 indicated impaired mitochondrial formation in *Dnah3* KO mice compared to WT mice. Red, SLC25A4; green, α-Tubulin; blue, DAPI; scale bars, 5 µm.

Moreover, subsequent computer-assisted sperm analysis (CASA) also showed that sperm isolated from the cauda epididymis were slightly decreased, and nearly all sperm were completely immobile (**Table 3**, **Movie S1 and Movie S2**). Papanicolaou staining and SEM analysis revealed morphological defects in partial spermatozoa from *Dnah3* KO mice, including coiled, bent, and irregular flagella, as well as aberrant heads and acephalic spermatozoa (**Figure S7A and B**).

**Table 3.** Semen analysis using CASA in the *Dnah3* KO mice.

|  | WT | KO | P<sup>b</sup> value |
| --- | --- | --- | --- |
| **Semen parameters** |  |  |  |
| Sperm concentration (10<sup>6</sup>/ml) <sup>a</sup> | 112.32 ± 18.26 | 105.17 ± 11.15 | 0.059 |
| Motility (%) <sup>b</sup> | 71.56 ± 3.97 | <u>4.37 ± 1.15</u> | <0.01 |
| Progressive motility (%) <sup>b</sup> | 60.36 ± 4.32 | <u>4.37 ± 1.15</u> | <0.01 |
| **Sperm locomotion parameters** |  |  |  |
| Curvilinear velocity (VCL) (μm/s) <sup>b</sup> | 67.54 ± 6.79 | <u>9.07 ± 1.22</u> | <0.01 |
| Straight-line velocity (VSL) (μm/s) <sup>b</sup> | 28.91 ± 4.86 | <u>2.68 ± 0.52</u> | <0.01 |
| Average path velocity (VAP) (μm/s) <sup>b</sup> | 39.02 ± 5.31 | <u>3.85 ± 0.82</u> | <0.01 |
| Amplitude of lateral head displacement (ALH) (μm) <sup>b</sup> | 0.71 ± 0.03 | <u>0.13 ± 0.04</u> | <0.01 |
| Linearity (LIN) <sup>b</sup> | 0.43 ± 0.07 | <u>0.30 ± 0.02</u> | 0.037 |
| Wobble (WOB, = VAP/VCL) <sup>b</sup> | 0.58 ± 0.06 | <u>0.42 ± 0.05</u> | 0.024 |
| Straightness (STR, = VSL/VAP) | 0.74 ± 0.22 | 0.70 ± 0.13 | 0.80 |
| Beat-cross frequency (BCF) (Hz) <sup>b</sup> | 4.86 ± 0.12 | <u>0.73 ± 0.08</u> | <0.01 |

<sup>a</sup> Epididymides and vas deferens.
<sup>b</sup> A significant difference, two-sided student's t-test. n = 3 biologically independent WT mice or KO mice.

TEM was further utilized to evaluate the sperm flagellar ultrastructure of *Dnah3* KO mice. There were no obvious abnormalities of “9 + 2” microtube arrangement in most sperm from the *Dnah3* KO mice when compared to WT mice (**Figure 4B**, **Figure S7C**). However, in contrast to the clear display of an IDA and an ODA on the A-tube of each microtubule doublet in the sperm flagella of WT mice, the sperm flagella of *Dnah3* KO mice exhibited an absence of almost all the IDAs (**Figure 4B**, **Figure S7D**). In addition, the disrupted mitochondria of spermatozoa from *Dnah3* KO mice were also observed under TEM, as manifested by the dilated intermembrane spaces and dissolved mitochondrial material (**Figure 4B and C**, **Figure S7E**). We next performed immunofluorescence staining to label SLC25A4, which is responsible for the exchange of ATP and ADP across the mitochondrial inner membrane. Strikingly, compared to the bright fluorescence signals in the midpiece of WT sperm, the signals *Dnah3* KO were significantly diminished (**Figure 4D**), indicating impaired mitochondrial function. Collectively, DNAH3 is essential for spermatogenesis, and its deficiency seriously damages the sperm motility and IDAs in both humans and mice.

### DNAH3 deficiency impairs IDAs related to the reduction of IDA-associated proteins

Considering the disrupted IDAs revealed by TEM analysis in both our patients and *Dnah3* KO mice, we speculated whether the defective IDAs were attributed to the decreased expression of the key IDA-associated proteins. The immunofluorescence data showed that DNAH1/DNAH6 and DNALI1, corresponding to the heavy and light intermediate chains of the IDAs (16), respectively, were almost invisible along the sperm flagella of the patients when compared to control (**Figure 5A-C**). Consistent results were obtained in our subsequent western blotting analysis of sperm lysates from the patients (**Figure 5D-F**), indicating that DNAH3 may manipulate the assembly of IDA through regulating the expression of IDA-associated proteins. In contrast, DNAH8/DNAH17 and DNAI1, corresponding to the heavy and intermediate chains of ODAs (25), were readily detectable in the patients’ sperm flagella and were comparable to the control (**Figure S8A-C**), suggesting that DNAH3 may not regulate the expression of ODA-associated proteins. We also performed immunofluorescence staining and western blotting analysis of DNAH1, DNAH6, DNALI1, DNAH8, DNAH17 and DNAI1 on sperm from *Dnah3* KO mice, and the results observed were consistent with those of the patients (**Figure 6A-F**, **Figure S9A-C**). These findings suggested that other IDA-associated proteins might be downstream effectors of DNAH3, which needs more future research.

**Figure 5.**
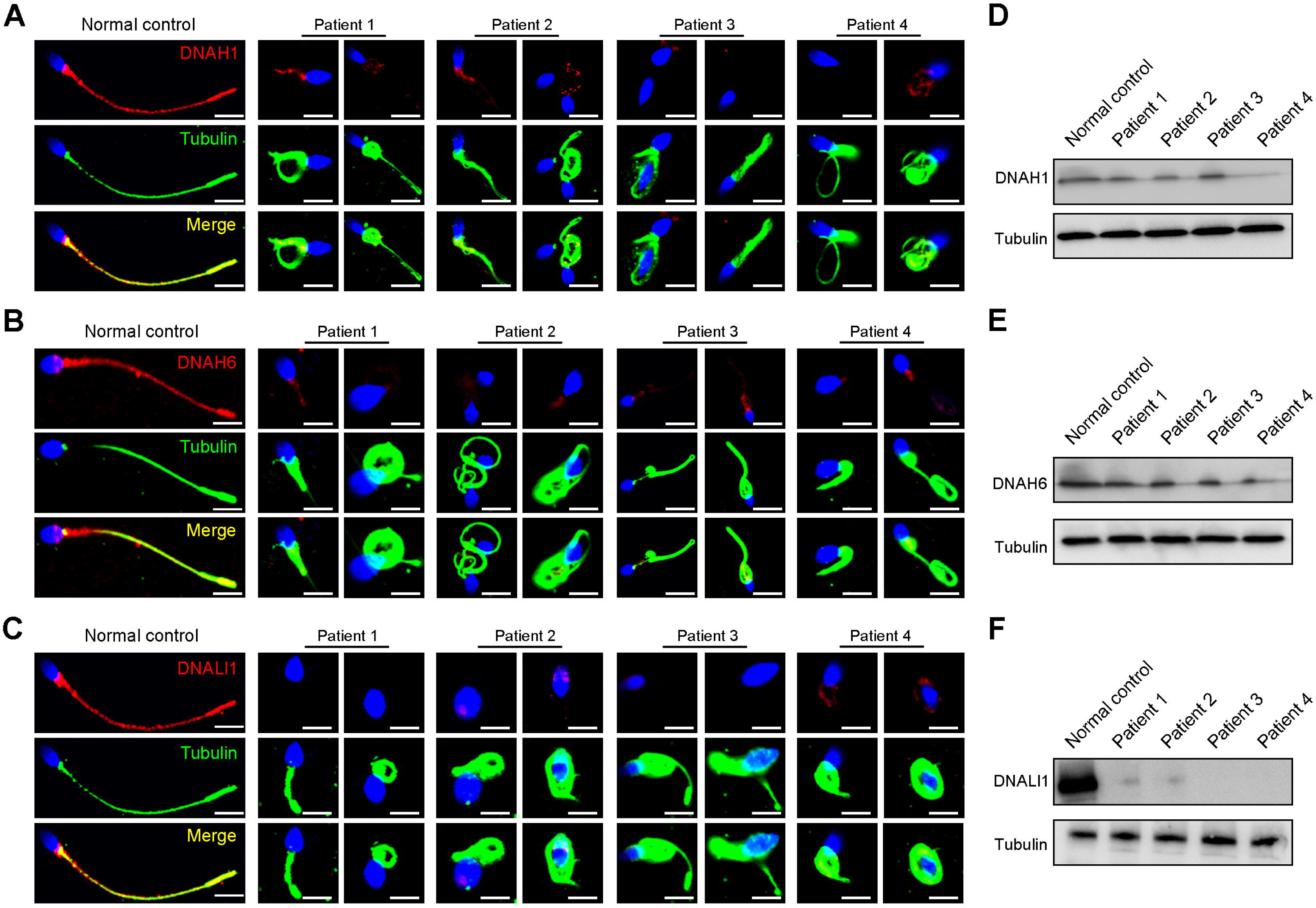
Immunofluorescence staining and western blotting analysis of IDA-associated proteins in spermatozoa obtained from normal control and patients with *DNAH3* variants. (**A** – **C**) Immunofluorescence staining of DNAH1 (A), DNAH6 (B) and DNALI1 (C) in spermatozoa from patients and normal controls. Red, DNAH1 in (A), DNAH6 in (B), DNALI1 in (C); green, α-Tubulin; blue, DAPI; scale bars, 5 μm. (**D** – **F**) Western blotting analysis of DNAH1(D), DNAH6 (E), DNALI1 (F) in sperm lysates from the patients and normal control.

**Figure 6.**
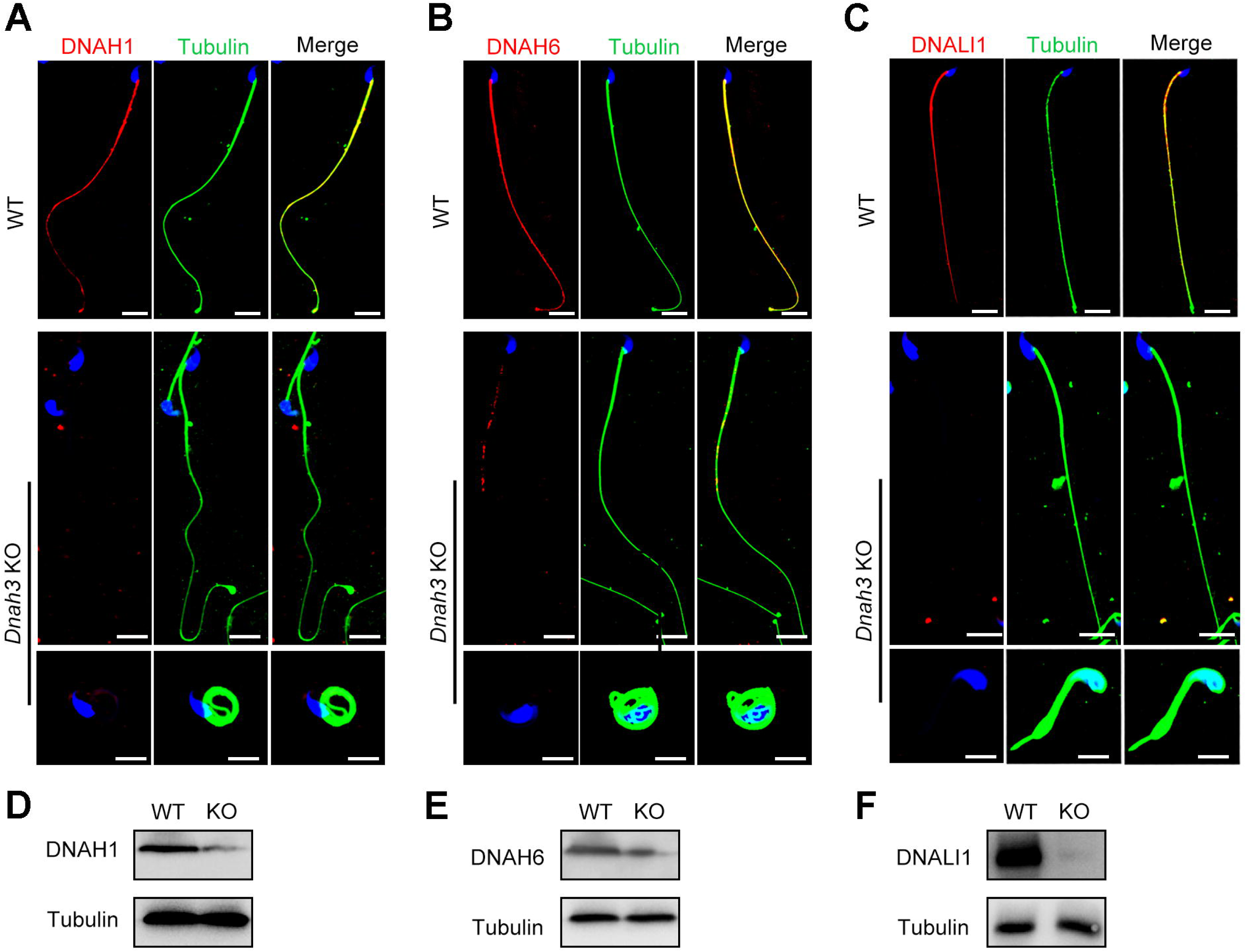
Immunofluorescence staining and western blotting analysis of IDA-associated proteins in spermatozoa from WT and *Dnah3* KO mice. (**A** – **C**) Immunofluorescence staining of DNAH1 (A), DNAH6 (B) and DNALI1 (C) in spermatozoa from *Dnah3* KO and WT mice. Red, DNAH1 in (A), DNAH6 in (B), DNALI1 in (C); green, α-Tubulin; blue, DAPI; scale bars, 5 μm. (**D** – **F)** Western blotting analysis of DNAH1(D), DNAH6 (E) and DNALI1 (F) in spermatozoa lysates from *Dnah3* KO and WT mice.

### ICSI treatment of humans with *DNAH3* variants and *Dnah3* KO mice

ICSI treatment has been reported to be effective in asthenoteratozoospermia-associated infertility (34, 35). ICSI cycles were attempted for the patients after written informed consent was obtained. The female partners all had normal basal hormone levels and underwent a long gonadotrophin-releasing hormone agonist protocol (**Table 4**). The wife of Patient 1 underwent one ICSI attempt. A total of 21 metaphase II (MII) oocytes were retrieved and microinjected, of which 17 oocytes were successfully fertilized (17/21, 80.95%) and cleaved (17/17, 100%). Thirteen Day 3 (D3) embryos were formed, six of which developed into blastocysts (8/13, 61.54%) after standard embryo culture. Two blastocysts were transferred, one of which was implanted. She eventually achieved clinical pregnancy, and the pregnancy is ongoing (**Table 4**). The partner of Patient 2 underwent two ICSI attempts. In her first ICSI attempt, six MII oocytes were retrieved, of which three were fertilized (3/6, 50%) and cleaved (3/3, 100%). After standard embryo culture, two D3 embryos were formed and transferred. However, this ICSI failed because no embryos were implanted. In her second ICSI attempt, all five MII oocytes were fertilized and cleaved (5/5, 100%). Five D3 embryos were obtained, of which two were transferred, but no embryos were implanted. The remaining three D3 embryos were cultured continuously, and two available blastocysts were formed and kept to be transferred in the future (**Table 4**). The partner of Patient 3 underwent one ICSI attempt. Of the 20 MII oocytes retrieved, 19 oocytes were fertilized (19/20, 95.0%) and cleaved (19/19, 100%). Fifteen D3 embryos were obtained, and 10 developed into available blastocysts (10/15, 66.7%). One blastocyst was transferred and implanted. She achieved clinical pregnancy, and the pregnancy is ongoing (**Table 4**). The wife of Patient 4 underwent four failed ICSI attempts. In her first two ICSI attempts, 13 and 12 MII oocytes were retrieved, of which five (5/12, 41.67%) and six (6/13, 46.15%), respectively, were fertilized and cleaved. Two available D3 embryos were obtained and transferred in both ICSI attempts, but no embryos were implanted. In her third ICSI attempt, of the eight MII oocytes retrieved, four (4/8, 50%) were fertilized and cleaved (4/4, 100%). However, no available D3 embryos were acquired. In her last ICSI attempt, seven MII oocytes were retrieved, of which three were fertilized (3/7, 42.68%) and two were cleaved (2/3, 66.7%), but no available D3 embryos were formed (**Table 4**). The vivid embryonic development of the partner of Patient 1 and Patient 3 after ICSI treatment was shown in **Figure 7A**.

**Figure 7.**
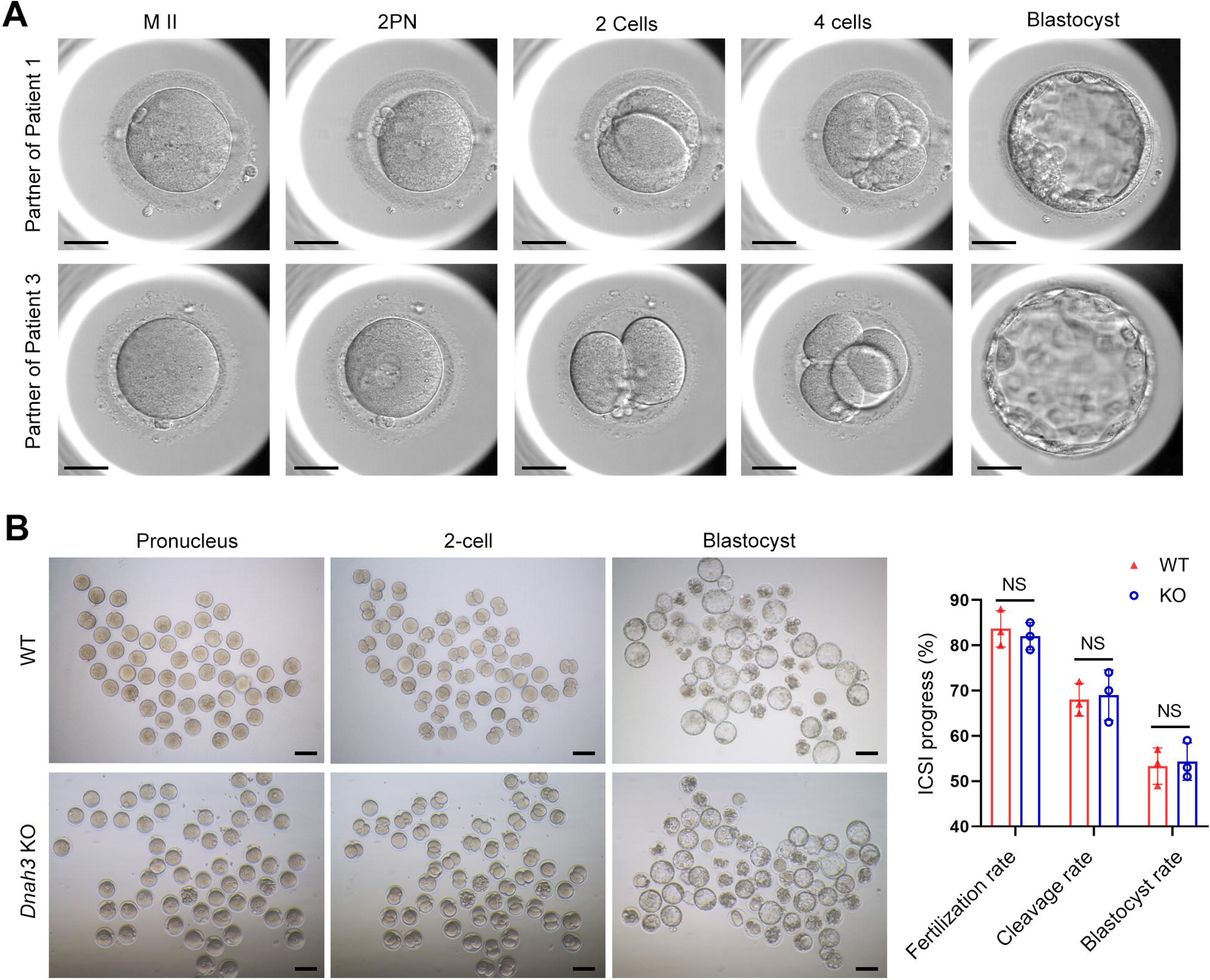
ICSI outcomes of *DNAH3*-deficient patients and *Dnah3* KO mice. (**A**) The embryonic development of Patient 1 and Patient 3 after ICSI treatment. MII, metaphase II; PN, pronucleus; scale bars, 40 μm. (**B**) There was no difference in the fertilization rate or 2-cell and blastocyst embryo formation rates between the *Dnah3* KO and WT groups (n = three biologically independent WT mice or KO mice; Student’s *t* test; NS, no significance; error bars, s.e.m.).

**Table 4.**
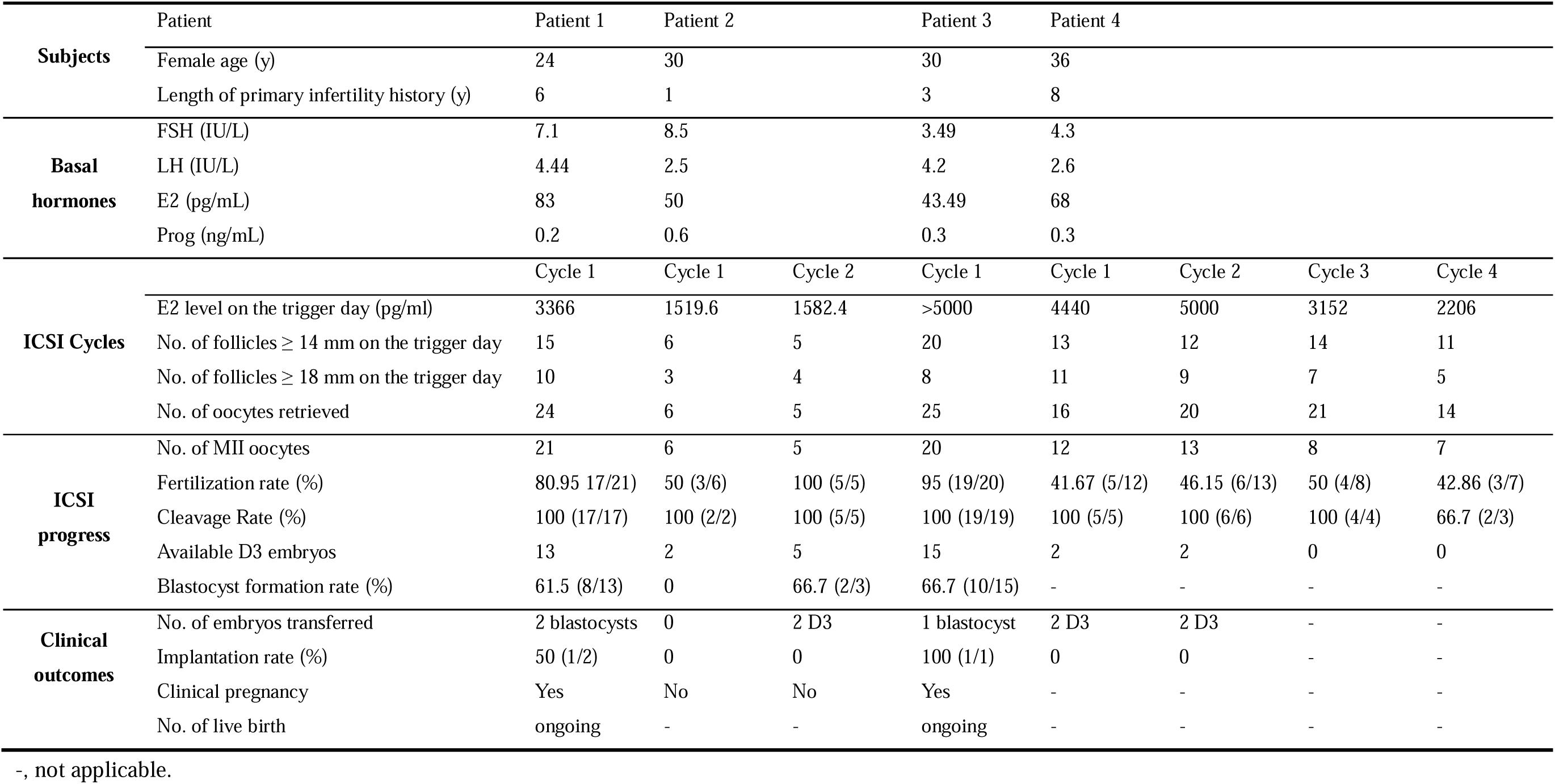
Outcomes of ICSI treatment in the patients with *DNAH3* mutations.

We also carried out ICSI treatment on *Dnah3* KO male mice. Strikingly, favorable outcomes of ICSI were obtained in *Dnah3* KO male mice. After injection of spermatozoa from *Dnah3* KO male mice, pronuclei were observed in most embryos in both the KO and WT groups, indicating a normal fertilization rate (**Figure 7B**). There was no difference in the percentage of 2-cell and blastocyst-stage embryos between the KO and WT groups (**Figure 7B**). Collectively, we observed successful ICSI outcomes in two out of four DNAH3-deficient patients and *Dnah3* KO male mice and therefore suggested ICSI as an optional treatment for infertile men harboring biallelic pathogenic variants in *DNAH3*, and the additional female risk factors for infertility should not be excluded in the failed patients.

## Discussion

In the present study, we identified pathogenic variants in *DNAH3* in unrelated infertile men with asthenoteratozoospermia. These variations resulted in the almost absence of DNAH3 and sharply decreased the expression of other IDA-associated proteins, including DNAH1, DNAH6 and DNALI1. Combined with similar findings in *Dnah3* KO mice, we demonstrated that DNAH3 is fundamental for male fertility. Moreover, we suggest that ICSI might be a favorable treatment for male infertility caused by DNAH3 deficiency. Our findings identify a function for DNAH3 in male reproduction in humans and mice and may provide a new view on the clinical practice of male infertility.

Recently, Meng et al. reported *DNAH3* mutations in asthenoteratozoospermia affected patients, revealing multiple morphological defects in sperm tail (36). Moreover, ultrastructural abnormalities of the flagellar axoneme in the patients were evident in these patients, characterized by a disrupted ’9+2’ arrangement and the notable absence of IDAs (36). Additionally, they generated *Dnah3* KO mice, which were infertile and exhibited moderate morphological abnormalities (36). While the ’9+2’ microtubule arrangement in the flagella of their *Dnah3* KO mice remained intact, the IDAs on the microtubules were partially absent (36). In our study, we observed similar phenotypic differences between *DNAH3*-deficient patients and *Dnah3* KO mice. Both studies suggest that *DNAH3* may play crucial yet distinct roles in human and mouse male reproduction.

However, there are notable differences between the two studies. Firstly, the phenotypes of *Dnah3* KO mice showed slight differences. Meng et al. generated two *Dnah3* KO mouse models (KO1 and KO2), and both of which exhibited significantly higher sperm motility and progressive motility than in our study (36), where nearly all sperm were completely immobile. Furthermore, their *Dnah3* KO2 mice displayed motility comparable to WT mice and retained partial fertility (36). We speculate that these differences may be attributed to variations in mouse genetic background or the presence of a truncated DNAH3 protein resulting from specific knockout strategies. Secondly, we conducted additional research and uncovered novel findings. We revealed that male infertility caused by *DNAH3* mutations follows an autosomal recessive inheritance pattern, as confirmed through Sanger sequencing of the patients’ families. We also discovered the dynamic expression and localization of DNAH3 during spermatogenesis in humans and mice through immunofluorescent staining. Initially, DNAH3 was expressed in the cytoplasm of spermatogonia and spermatocytes, and then it clearly transferred into the flagellum of early and late spermatids. We further found that DNAH3 deficiency had no impact on ciliary development in the oviduct or on oogenesis in mice, resulting in normal female fertility. Moreover, in the absence of DNAH3 in both humans and mice, the expression of IDA-associated proteins, including DNAH1, DNAH6 and DNALI1, was decreased, while the expression of ODA-associated proteins remained unaffected, indicating that DNAH3 is involved in sperm axonemal development, specifically through its role in the assembly of IDAs. Collectively, our study corroborates the findings of Meng et al., and provides additional unique insights, comprehensively elucidating the critical role of DNAH3 in human and mouse spermatogenesis.

Primary ciliary dyskinesia (PCD, MIM: 244400) is a genetic disorder affecting at least one in 7554 individuals (37). The most common symptoms of PCD are recurrent infections in airways due to malfunction of the motile cilia that are responsible for mucus clearance (38). It has been suggested that male infertility associated with sperm defects is highly prevalent (up to 75%) among individuals with PCD (39). Axonemal defects caused by variants within *DNAH* family members, including *DNAH5*, *DNAH6*, *DNAH7*, *DNAH9* and *DNAH11,* are causative factors for PCD (40–42). Moreover, deficiency in these PCD-causing *DNAH*s has also been associated with male infertility (9, 14, 20, 21, 43–45). Additionally, other *DNAH*s, such as *DNAH1*, *DNAH2*, *DNAH8*, *DNAH10*, *DNAH12* and *DNAH17,* are suggested to be pathogenic genes of isolated male infertility (18, 19, 22, 24, 25). These phenotype–genotype correlations may be attributed to the fact that ciliary and flagellar axonemes have cell type-specific or cell type-enriched DNAHs (46). DNAH3 resides in the IDA and is expressed in testis and ciliary tissues, including the lung, brain, eye, and oviduct. However, despite its presence in these tissues, the relationship between deficient DNAH3 and disease is unclear to date. Intriguingly, in our study, none of the patients with *DNAH3* deficiency reported experiencing any of the principal symptoms associated with PCD. Additionally, our *Dnah3* KO mice exhibited normal ciliary development in the lung, brain, eye, and oviduct. Similarly, Meng et al. did not mention any PCD symptoms in their *DNAH3*-deficient patients, and their *Dnah3* KO mice also demonstrated normal ciliary morphology in the trachea and brain (36). These combined observations suggest that DNAH3 may play a more important role in sperm flagellar development than in other motile cilia functions. Given that DNAH3 is expressed in ciliary tissues, its role in these tissues remains intriguing and could be elucidated through sequencing of larger cohorts of individuals with PCD.

ICSI has been an efficient treatment for male infertility (47, 48). However, the outcomes of ICSI for male infertility caused by variants in different *DNAH* genes are variable. It has been demonstrated that infertile males with variants in *DNAH1*, *DNAH2*, *DNAH7*, and *DNAH8* have a favorable prognosis (22, 49–53), while patients with variants in *DNAH17* have poor outcomes after ICSI treatment (25, 54). Meanwhile, the ICSI outcomes in male infertility caused by *DNAH6* variants may depend on the specific mutation or be controversial (20, 55, 56). The patients with *DNAH3* mutations in our study experienced different clinical outcomes of ICSI treatment. The partners of Patient 1 and Patient 3 achieved clinical pregnancy. The wives of Patient 2 and Patient 4 obtained favorable fertilization and cleavage rates but experienced no clinical pregnancy due to the nonimplantation of the transferred embryos. Remarkably, despite the diverse variants within *DNAH3* observed in the four patients, all variants led to a complete absence of DNAH3 expression. Additionally, we did not identify any pathogenic variants that associated with fertilization failure and early embryonic development in the two patients with failed ICSI outcomes. Therefore, these different ICSI outcomes might be attributed to additional unexplained factors from the female partners. Importantly, in the study from Meng et al., one patient carrying *DNAH3* variants received ICSI treatment, and the partner obtained clinical pregnancy (36). Combined with the successful ICSI outcomes observed in *Dnah3* KO mice, we suggest ICSI as an optimized treatment for infertile men carrying variants in *DNAH3*. More cases are needed to precisely estimate the prevalence of *DNAH3* mutations and determine a prognosis for ICSI treatments.

In conclusion, our study revealed an unexplored role of DNAH3 in male reproduction in humans and mice, suggesting *DNAH3* as a novel causative gene for human asthenoteratozoospermia. Moreover, ICSI is as an optimized treatment for infertile men with *DNAH3* variants. This study expands our knowledge of the relationship between DNAH proteins and disease, facilitating genetic counseling and clinical treatment of male infertility in the future.

## Methods

### Human subjects

Four unrelated Han Chinese infertile men and their family members were recruited from West China Second University Hospital of Sichuan University and Women and Children’s Hospital of Chongqing Medical University. All patients exhibited a normal karyotype (46 XY) without deletion of the azoospermia factor (AZF) region in the Y-chromosome. All of the participants were provided informed consent, and the study was approved by the ethics committee of West China Second University Hospital and The First Affiliated Hospital of Chongqing Medical University.

### Genetic analysis

Peripheral blood samples were obtained from the subjects to extract genomic DNA using a DNA purification kit (TIANGEN, DP304). For WES, 1 μg of genomic DNA was utilized for exon capture using the Agilent SureSelect Human All Exon V6 Kit and sequenced on the Illumina HiSeq X system (150-bp read length). The quality of WES, including clean reads, sequencing depth, sequencing coverage, and mapping quality are listed in **Table S1**. The variants identified through WES were annotated and filtered using Exomiser. Next, the variants were screened to obtain candidate variants based on the following criteria: (1) the allele frequency in the East Asian population was less than 1% in any database, including the ExAC Browser, gnomAD, and the 1000 Genomes Project; (2) the variants affected coding exons or canonical splice sites; (3) the variants were predicted to be possibly pathogenic or damaging. The remain genes were then analyzed using the Human Protein Atlas (HPA) database (https://www.proteinatlas.org/) and Mouse Genome Informatics (MGI) database (https://informatics.jax.org/) to access their expression in human and mouse testis. Additionally, OMIM database (https://www.omim.org/) and relevant literature were used to understand their relationship with human infertility. Given the assumption of a recessive inheritance pattern, monoallelic variants were excluded from consideration. The remained candidate pathogenic variants were verified by Sanger sequencing on DNA from the patients’ families. The primer pairs used for PCR amplification are listed in **Table S2**.

### Electron microscopy

For scanning electron microscopy (SEM), sperm samples were fixed in glutaraldehyde (2.5%, w/v) and dehydrated using an ethanol gradient (30, 50, 75, 85, 95, and 100% ethanol). The samples were dried using a CO_2_ critical-point dryer (Eiko HCP-2, Hitachi) and observed under SEM (S-3400, Hitachi).

For transmission electron microscopy (TEM), sperm samples were fixed in glutaraldehyde (3%, w/v) and osmium tetroxide (1%, w/v) and dehydrated with an ethanol gradient. The samples were embedded in Epon 812. Ultrathin sections were stained with uranyl acetate and lead citrate and analyzed under TEM (Tecnai G2 F20).

### STA-PUT velocity sedimentation

Single testicular cells from obstructive azoospermia and 8-week-old C57BL male mice were obtained using the STA-PUT velocity sedimentation method as described previously (57, 58). In brief, total spermatogenic cells were harvested by digesting seminiferous tubules with collagenase (Invitrogen, 17100017), trypsin (Sigma, T4799) and DNase (Promega, M6101) for 15 min each at 37 °C. Cells were diluted in bovine serum albumin (BSA, 3%, w/v) and filtered through an 80 mm mesh to remove fragments. Then, the cells were resuspended in BSA (3%, w/v) and loaded into an STA-PUT velocity sedimentation cell separator (ProScience) to obtain germ cells at different stages.

### RNA isolation and quantitative PCR (qPCR)

Total RNA of mouse tissues was extracted using TRIzol reagent (Invitrogen,15596026,) and reverse-transcribed using the 1st Strand cDNA Synthesis Kit (Yeasen, HB210629) according to the manufacturer’s instructions. qPCR was carried out on an iCycler RTLPCR Detection System (Bio-Rad Laboratories) using SYBR Green qPCR Master Mix (Bimake, B21202). Primer sequences are listed in **Table S2**.

### Immunofluorescence staining

Sperm samples were fixed in paraformaldehyde (4%, w/v), permeabilized with Triton X-100 (0.3% v/v) and blocked with BSA (3%, w/v) at room temperature. Samples were incubated with primary antibodies, including DNAH1 (Cusabio, CSB-PA878961LA01HU, 1:100), DNAH3 (Cusabio, CSB-PA823461LA01HU, 1:100), DNAH6 (Proteintech, 18080-1-AP, RRID: AB_2878493, 1:50), DNAH8 (Atlas, HPA028447, RRID: AB_10599600, 1:200), DNAH17 (Proteintech, 24488-1-AP, RRID: AB_2879568, 1:50), DNAI1 (Proteintech, 12756-1-AP, RRID: AB_10643244, 1:50), DNALI1 (Proteintech, 17601-1-AP, RRID: AB_2095372, 1:50), TOM20 (Proteintech, 11802-1-AP, RRID: AB_2207530, 1:50), SLC25A4 (Signalway, 32484, RRID: AB_2941094, 1:100), lectin PNA (Invitrogen, L-32460, 1:50) and alpha tubulin (Abcam, ab7291, RRID: AB_2241126, 1:500), overnight at 4 °C. The next day, the samples were washed and incubated with the secondary antibody Alexa Fluor 488 (Invitrogen, A11008, RRID: AB_143165, 1:1000) or Alexa Fluor 594 (Invitrogen, A11005, RRID: AB_141372, 1:1000), and the nuclei were labeled with 4′,6-diamidino-2-phenylindole (DAPI, SigmaLAldrich, D9542). Image capture was performed by a laser scanning confocal microscope (Olympus, FV3000).

For staining of mouse tissues, samples were first fixed in paraformaldehyde (4%, w/v) and dehydrated with an ethanol gradient. Then, the samples were embedded in paraffin and sliced into 5-μm sections. After deparaffinization and rehydration, sections were processed with 3% hydrogen peroxide and incubated in sodium citrate for antigen repair. Subsequently, sections were blocked with goat serum and incubated with primary antibodies against DNAH3 (Cusabio, CSB-PA823461LA01HU, 1:100) or ac-Tubulin (Abcam, ab24610, RRID: AB_448182, 1:500) at 4 °C overnight. The next day, the sections were incubated with the secondary antibody Alexa Fluor 488, followed by labeling the nuclei with DAPI. Image capture was performed using a fluorescence microscope (Zeiss, Ax10).

### Western Blotting

Sperm samples were lysed in RIPA buffer (Beyotime, P0013B) to extract the total protein. For analysis of DNALI1, the protein samples were mixed with SDS loading buffer (P0015, Beyotime, China), boiled at 95 °C for 5 minutes, and separated by 12.5% SDS-PAGE. For analysis of DNAH1, DNAH3, DNAH6, the protein samples were mixed with NuPAGE™ LDS sample buffer (Invitrogen, NP0007), denatured at 70°C for 10 minutes, and separated by 3–8% NuPAGE™ Tris-Acetate gels (EA0375BOX, Invitrogen). Then the resolved proteins were transferred to 0.45 μm PVDF membranes (Merck Millipore, IPVH00010). The membranes were blocked, incubated with primary antibodies, including DNALI1 (Proteintech, 17601-1-AP, RRID: AB_2095372, 1:150), DNAH3 (Cusabio, CSB-PA823461LA01HU, 1:200), DNAH6 (Proteintech, 18080-1-AP, RRID: AB_2878493, 1:150) and alpha tubulin (Proteintech, 11224-1-AP, RRID: AB_2210206, 1:1000) at 4 L overnight. The following day, membranes were washed and incubated with HRP-conjugated secondary antibody (Proteintech, SA00001-2, RRID: AB_2722564, 1:5000). Protein bands were visualized using enhanced chemiluminescence reagents (Millipore, WBKLS0500).

### Histology hematoxylin-eosin (H&E) staining

Tissue samples from mice were fixed with 4% paraformaldehyde (w/v) overnight. Following dehydration by ethanol, the samples were embedded in paraffin and sliced into 5-μm sections. The sections were stained with hematoxylin and eosin and observed under a microscope (Zeiss, Axio Imager 2).

### Generation of the *Dnah3* KO mouse model

Animal experiments in this study were approved by the Experimental Animal Management and Ethics Committee of West China Second University Hospital, Sichuan University, and complied with the Animal Care and Use Committee of Sichuan University. A *Dnah3* knockout mouse model was generated by the CRISPRLCas9 system. Briefly, Cas9 and signal-guide RNAs (5’-GTATCAAGTGGATGTAAACC-3’) were transcribed using T7 RNA polymerase in vitro and comicroinjected into the cytoplasm of single-cell C57BL/6J mouse embryos to generate frameshift mutations by nonhomologous recombination through introduction of a 1 bp insertion in exon 13. Then, the embryos were cultured and transferred into the oviducts of pseudopregnant female mice at 0.5 days post-coitum. A mutation of *Dnah3* in the founder mouse and their offspring was confirmed using PCR and Sanger sequencing. The primers used for the generation of animal models are listed in **Table S2.**

### Intracytoplasmic sperm injection (ICSI)

ICSI was carried out using standard techniques. In brief, one-month-old female KM mice were injected with 5 IU of equine chorionic gonadotropin (eCG) (ProSpec, HOR-272) to induce superovulation. Metaphase II-arrested (MII) oocytes were acquired through another injection of 5 IU human chorionic gonadotropin after 48 hours. MII oocytes were incubated with Chatot-Ziomek-Bavister medium (Easycheck, M2750) at 37.5 °C and 5% CO_2_ until use. Mouse cauda epididymal spermatozoa were incubated in human tubal fluid (HTF) medium (Easycheck, M1150) and then frozen and thawed repeatedly to remove sperm tails. For ICSI, a single sperm head was microinjected into an MII oocyte by using a NIKON inverted microscope and a Piezo (PrimeTech, Osaka, Japan) in Whitten’s-HEPES medium containing 0.01% polyvinyl alcohol (Gibco,12360-038) and cytochalasin B (3.5 g/ml; SigmaLAldrich, C-6762). The successfully injected oocytes were transferred into G1-Plus medium (Vitrolife, 10132) and incubated at 37.5 °C and 5% CO_2_. The animal experiments were approved by the Experimental Animal Management and Ethics Committee of West China Second University Hospital, Sichuan University.

### Statistical analysis

Prism (version 8.4.0, GraphPad, Boston, MA, USA) and SPSS (version 18.0, IBM Corporation, Armonk, NY, USA) were used to perform statistical analyses. All data are presented as the means ± SEMs. Data from two groups were compared using an unpaired, parametric, two-sided Student’s *t* test, and a *p* value less than 0.05 was considered statistically significant.

## Ethics approval

This study was approved by Ethical Review Board of West China Second University Hospital, Sichuan University. Informed consent was obtained from each participate in this study before taking part.

## Data availability

The published article includes all datasets generated or analyzed during this study. The whole exome-sequencing data were deposited in the National Genomics Data Center (NGDC) (https://ngdc.cncb.ac.cn/, accession number: HRA007467).

## Competing interests

The authors declare that they have no conflict of interest.

## Author contributions

**Xiang Wang**: Data curation; formal analysis; investigation; methodology; writing – original draft. **Gan Shen**: Data curation; formal analysis; **Yihong Yang**: Resources; investigation; methodology. **Chuan Jiang**: Data curation; formal analysis. **Tiechao Ruan**: Formal analysis; methodology. **Xue Yang**: Validation; investigation. **Liangchai Zhuo**: Investigation; **Yingteng Zhang**: Investigation, methodology. **Yangdi Ou**: Investigation. **Xinya zhao**: Investigation. **Shunhua Long**: Methodology. **Xiangrong Tang**: Investigation. **Tingting Lin**: Conceptualization; funding acquisition; project administration. **Ying Shen**: Conceptualization; supervision; project administration; writing – review and editing.

## Acknowledgements

The authors thank the patient and his family members for their voluntary participation. We are grateful to Guiping Yuan from Analytical and Testing Center of Sichuan University and Yan Liang from Research Core Facility of West China Hospital, Sichuan University for their help with TEM images and preparing histology slides. This work was supported by National Natural Science Foundation of China (82301807).

## Supporting information

**Figure S1. Statistics analysis of aberrant sperm morphology and axonemal ultrastructure observed in *DNAH3*-deficient patients. (A, B)** The percentage distribution (A) and histogram (B) of various flagellar morphology in the normal control and patients. (**C**) The percentage of ultrastructure in different cross-sections of sperm from the normal control and patients (n = three biologically independent WT mice or KO mice; error bars, s.e.m.).

**Figure S2. The expression of DNAH3 in mouse testis.** (**A**) qPCR analysis revealed that *Dnah3* was highly expressed in the mouse testis. (**B**) qPCR analysis showed that *Dnah3* expression was significantly elevated beginning on postnatal Day 12, peaked at postnatal Day 30, and maintained a stable expression level thereafter.

**Figure S3. DNAH3 is expressed during spermatogenesis in mice and humans.** (**A**) Immunofluorescence staining of DNAH3 in isolated mouse germ cells. Pink, PNA; green, DNAH3; blue, DAPI; scale bars, 5 μm. (**B**) Immunofluorescence staining of DNAH3 in isolated human germ cells. Pink, PNA; green, DNAH3; blue, DAPI; scale bars, 5 μm.

**Figure S4. Generation of *Dnah3* KO mice.** (**A**) Schematic illustration of the strategy for the generation of *Dnah3* KO mice. (**B, C**) PCR sequencing (B) and qPCR (C) were used to confirm the genotype and KO efficiency (n = three biologically independent WT mice or KO mice; Student’s *t* test; *, P<0.05; error bars, s.e.m.). (**D**) Immunofluorescence staining of DNAH3 in testis of *Dnah3* KO mice and WT mice. Green, DNAH3; blue, DAPI; scale bars, 75 μm. (**E**) Immunofluorescence staining of DNAH3 in spermatozoa isolated from the cauda epididymis of *Dnah3* KO mice and WT mice. Red, DNAH3; green, α-Tubulin; blue, DAPI; scale bars, 5 μm.

**Figure S5. Ciliary development of *Dnah3* KO mice.** (**A**) H&E staining of lung, brain, eye, and oviduct from *Dnah3* KO mice and WT mice. Scale bars, 100 μm. (**B**) Analysis of ciliary development in the lung, brain, eye, and oviduct from *Dnah3* KO mice and WT mice by using immunofluorescence staining. Green, Ac-Tubulin; blue, DAPI; scale bars, 20 μm.

**Figure S6. Fertility of *Dnah3* KO mice.** (**A**) H&E staining of ovary tissue sections from 8-week-old *Dnah3* KO female mice and WT female mice. Scale bars, 75 μm (n = three biologically independent WT mice or KO mice). (**B**) Sizes of the testis and epididymis of the 8-week-old *Dnah3* KO and WT mice (n = three biologically independent WT mice or KO mice; Student’s *t* test; NS, no significance; error bars, s.e.m.).

**Figure S7. Morphology and ultrastructure of sperm isolated from *Dnah3* KO mice. (A, B)** Papanicolaou staining (A), and SEM analysis (B) revealed morphological defects in partial spermatozoa from *Dnah3* KO mice compared to WT mice. Scale bars in (A), 5 μm; scale bars in (B), 2.5 μm. (n = three biologically independent WT mice or KO mice; Student’s *t* test; error bars, s.e.m.). **(C)** The percentage of aberrant axonemal arrangement in different cross-sections of sperm from WT mice and *Dnah3* KO mice. (n = three biologically independent WT mice or KO mice; error bars, s.e.m.). (**D**) The percentage of microtubule doublets that presented IDAs in WT mice and *Dnah3* KO mice. (n = three biologically independent WT mice or KO mice; Student’s *t* test; error bars, s.e.m.). (**E**) Statistics of malformed mitochondria in the midpiece of sperm from WT mice and *Dnah3* KO mice. (n = three biologically independent WT mice or KO mice; Student’s *t* test; error bars, s.e.m.).

**Figure S8. Immunofluorescence staining of ODA-associated proteins in spermatozoa obtained from variants within *DNAH3* patients**. (**A** – **C**) The expression of DNAH8 (A), DNAH17 (B) and DNAI1 (C) in spermatozoa of the patients was comparable to that in normal controls. Red, DNAH8 in (A), DNAH17 in (B), DNAI1 in (C); green, α-Tubulin; blue, DAPI; scale bars, 5 μm.

**Figure S9. Immunofluorescence staining of ODA-associated proteins in spermatozoa of *Dnah3* KO and WT mice**. (**A** – **C**) The expression of DNAH8 (A), DNAH17 (B) and DNAI1 (C) in spermatozoa from *Dnah3* KO mice was comparable to that in spermatozoa from WT mice. Red, DNAH8 in (A), DNAH17 in (B), DNAI1 in (C); green, α-Tubulin; blue, DAPI; scale bars, 5 μm.

**Table S1. Summary of whole exome sequencing and the candidate variants identified.**

**Table S2. Primer pairs used in the present study.**

**Movie S1. CASA of sperm from WT mice**. Sperm from the epididymis of WT mice were collected, incubated, and recorded under a phase-contrast microscope. A normal quantity and motility of sperm were observed in the WT mice (n = three biologically independent WT mice).

**Movie S2. CASA of sperm from *Dnah3* KO mice**. Epididymal sperm of *Dnah3* KO mice were collected, incubated in HTF medium at 37 °C for 10 minutes, and recorded under a phase-contrast microscope. The movie showed a significantly reduced motility of sperm from *Dnah3* KO (n = three biologically independent *Dnah3* mice).

## Notes

https://ngdc.cncb.ac.cn/

### Competing Interest Statement

The authors have declared no competing interest.

### Summary of Updates

We have updated the title of our study to: 'DNAH3 deficiency causes flagellar inner dynein arm loss and male infertility in humans and mice'. Furthermore, we have highlighted the differences between our study and the recent study on DNAH3 from Meng et al. in the 'Discussion' section.

## Reference

1. Cox CM, Thoma ME, Tchangalova N, Mburu G, Bornstein MJ, Johnson CL, Kiarie J. Infertility prevalence and the methods of estimation from 1990 to 2021: a systematic review and meta-analysis. Hum Reprod Open. 2022;2022(4):hoac051.

2. Eisenberg ML, Esteves SC, Lamb DJ, Hotaling JM, Giwercman A, Hwang K, Cheng YS. Male infertility. Nat Rev Dis Primers. 2023;9(1):49.

3. Agarwal A, Mulgund A, Hamada A, Chyatte MR. A unique view on male infertility around the globe. Reprod Biol Endocrinol. 2015;13:37.

4. Toure A, Martinez G, Kherraf ZE, Cazin C, Beurois J, Arnoult C, et al. The genetic architecture of morphological abnormalities of the sperm tail. Hum Genet. 2021;140(1):21–42.

5. Wang WL, Tu CF, Tan YQ. Insight on multiple morphological abnormalities of sperm flagella in male infertility: what is new? Asian J Androl. 2020;22(3):236–45.

6. Wang J, Wang W, Shen L, Zheng A, Meng Q, Li H, Yang S. Clinical detection, diagnosis and treatment of morphological abnormalities of sperm flagella: A review of literature. Front Genet. 2022;13:1034951.

7. Lu S, Gu Y, Wu Y, Yang S, Li C, Meng L, et al. Bi-allelic variants in human WDR63 cause male infertility via abnormal inner dynein arms assembly. Cell Discov. 2021;7(1):110.

8. Houston BJ, Riera-Escamilla A, Wyrwoll MJ, Salas-Huetos A, Xavier MJ, Nagirnaja L, et al. A systematic review of the validated monogenic causes of human male infertility: 2020 update and a discussion of emerging gene-disease relationships. Hum Reprod Update. 2021;28(1):15–29.

9. Jiao SY, Yang YH, Chen SR. Molecular genetics of infertility: loss-of-function mutations in humans and corresponding knockout/mutated mice. Hum Reprod Update. 2021;27(1):154–89.

10. Leung MR, Zeng J, Wang X, Roelofs MC, Huang W, Zenezini Chiozzi R, et al. Structural specializations of the sperm tail. Cell. 2023;186(13):2880–96 e17.

11. Burgess SA, Walker ML, Sakakibara H, Knight PJ, Oiwa K. Dynein structure and power stroke. Nature. 2003;421(6924):715-8.

12. Linck RW, Chemes H, Albertini DF. The axoneme: the propulsive engine of spermatozoa and cilia and associated ciliopathies leading to infertility. J Assist Reprod Genet. 2016;33(2):141–56.

13. Gunes S, Sengupta P, Henkel R, Alguraigari A, Sinigaglia MM, Kayal M, et al. Microtubular Dysfunction and Male Infertility. World J Mens Health. 2020;38(1):9–23.

14. Sironen A, Shoemark A, Patel M, Loebinger MR, Mitchison HM. Sperm defects in primary ciliary dyskinesia and related causes of male infertility. Cell Mol Life Sci. 2020;77(11):2029–48.

15. King SM. Axonemal Dynein Arms. Cold Spring Harb Perspect Biol. 2016;8(11).

16. Walton T, Gui M, Velkova S, Fassad MR, Hirst RA, Haarman E, et al. Axonemal structures reveal mechanoregulatory and disease mechanisms. Nature. 2023;618(7965):625–33.

17. Aprea I, Raidt J, Hoben IM, Loges NT, Nothe-Menchen T, Pennekamp P, et al. Defects in the cytoplasmic assembly of axonemal dynein arms cause morphological abnormalities and dysmotility in sperm cells leading to male infertility. PLoS Genet. 2021;17(2):e1009306.

18. Ben Khelifa M, Coutton C, Zouari R, Karaouzene T, Rendu J, Bidart M, et al. Mutations in DNAH1, which encodes an inner arm heavy chain dynein, lead to male infertility from multiple morphological abnormalities of the sperm flagella. Am J Hum Genet. 2014;94(1):95–104.

19. Hwang JY, Nawaz S, Choi J, Wang H, Hussain S, Nawaz M, et al. Genetic Defects in DNAH2 Underlie Male Infertility With Multiple Morphological Abnormalities of the Sperm Flagella in Humans and Mice. Front Cell Dev Biol. 2021;9:662903.

20. Tu C, Nie H, Meng L, Yuan S, He W, Luo A, et al. Identification of DNAH6 mutations in infertile men with multiple morphological abnormalities of the sperm flagella. Sci Rep. 2019;9(1):15864.

21. Gao Y, Liu L, Shen Q, Fu F, Xu C, Geng H, et al. Loss of function mutation in DNAH7 induces male infertility associated with abnormalities of the sperm flagella and mitochondria in human. Clin Genet. 2022;102(2):130–5.

22. Liu C, Miyata H, Gao Y, Sha Y, Tang S, Xu Z, et al. Bi-allelic DNAH8 Variants Lead to Multiple Morphological Abnormalities of the Sperm Flagella and Primary Male Infertility. Am J Hum Genet. 2020;107(2):330–41.

23. Tu C, Cong J, Zhang Q, He X, Zheng R, Yang X, et al. Bi-allelic mutations of DNAH10 cause primary male infertility with asthenoteratozoospermia in humans and mice. Am J Hum Genet. 2021;108(8):1466–77.

24. Li Y, Wang Y, Wen Y, Zhang T, Wang X, Jiang C, et al. Whole-exome sequencing of a cohort of infertile men reveals novel causative genes in teratozoospermia that are chiefly related to sperm head defects. Hum Reprod. 2021;37(1):152–77.

25. Whitfield M, Thomas L, Bequignon E, Schmitt A, Stouvenel L, Montantin G, et al. Mutations in DNAH17, Encoding a Sperm-Specific Axonemal Outer Dynein Arm Heavy Chain, Cause Isolated Male Infertility Due to Asthenozoospermia. Am J Hum Genet. 2019;105(1):198–212.

26. Chapelin C, Duriez B, Magnino F, Goossens M, Escudier E, Amselem S. Isolation of several human axonemal dynein heavy chain genes: genomic structure of the catalytic site, phylogenetic analysis and chromosomal assignment. FEBS Lett. 1997;412(2):325–30.

27. Karak S, Jacobs JS, Kittelmann M, Spalthoff C, Katana R, Sivan-Loukianova E, et al. Diverse Roles of Axonemal Dyneins in Drosophila Auditory Neuron Function and Mechanical Amplification in Hearing. Sci Rep. 2015;5:17085.

28. Modiba MC, Nephawe KA, Mdladla KH, Lu W, Mtileni B. Candidate Genes in Bull Semen Production Traits: An Information Approach Review. Vet Sci. 2022;9(4).

29. Hamdi Y, Boujemaa M, Ben Rekaya M, Ben Hamda C, Mighri N, El Benna H, et al. Family specific genetic predisposition to breast cancer: results from Tunisian whole exome sequenced breast cancer cases. J Transl Med. 2018;16(1):158.

30. Ng PC, Henikoff S. SIFT: Predicting amino acid changes that affect protein function. Nucleic Acids Res. 2003;31(13):3812–4.

31. Adzhubei IA, Schmidt S, Peshkin L, Ramensky VE, Gerasimova A, Bork P, et al. A method and server for predicting damaging missense mutations. Nat Methods. 2010;7(4):248–9.

32. Schwarz JM, Cooper DN, Schuelke M, Seelow D. MutationTaster2: mutation prediction for the deep-sequencing age. Nat Methods. 2014;11(4):361–2.

33. Rentzsch P, Witten D, Cooper GM, Shendure J, Kircher M. CADD: predicting the deleteriousness of variants throughout the human genome. Nucleic Acids Res. 2019;47(D1):D886–D94.

34. Tsai CC, Huang FJ, Wang LJ, Lin YJ, Kung FT, Hsieh CH, Lan KC. Clinical outcomes and development of children born after intracytoplasmic sperm injection (ICSI) using extracted testicular sperm or ejaculated extreme severe oligo-astheno-teratozoospermia sperm: a comparative study. Fertil Steril. 2011;96(3):567–71.

35. Colpi GM, Francavilla S, Haidl G, Link K, Behre HM, Goulis DG, et al. European Academy of Andrology guideline Management of oligo-astheno-teratozoospermia. Andrology. 2018;6(4):513–24.

36. Meng GQ, Wang Y, Luo C, Tan YM, Li Y, Tan C, et al. Bi-allelic variants in DNAH3 cause male infertility with asthenoteratozoospermia in humans and mice. Hum Reprod Open. 2024;2024(1):hoae003.

37. Hannah WB, Seifert BA, Truty R, Zariwala MA, Ameel K, Zhao Y, et al. The global prevalence and ethnic heterogeneity of primary ciliary dyskinesia gene variants: a genetic database analysis. Lancet Respir Med. 2022;10(5):459–68.

38. O’Connor MG, Mosquera R, Metjian H, Marmor M, Olivier KN, Shapiro AJ. Primary Ciliary Dyskinesia. CHEST Pulmonary. 2023;1(1):100004.

39. Vanaken GJ, Bassinet L, Boon M, Mani R, Honore I, Papon JF, et al. Infertility in an adult cohort with primary ciliary dyskinesia: phenotype-gene association. Eur Respir J. 2017;50(5).

40. Hornef N, Olbrich H, Horvath J, Zariwala MA, Fliegauf M, Loges NT, et al. DNAH5 mutations are a common cause of primary ciliary dyskinesia with outer dynein arm defects. Am J Respir Crit Care Med. 2006;174(2):120–6.

41. Guan Y, Yang H, Yao X, Xu H, Liu H, Tang X, et al. Clinical and Genetic Spectrum of Children With Primary Ciliary Dyskinesia in China. Chest. 2021;159(5):1768–81.

42. Peng B, Gao YH, Xie JQ, He XW, Wang CC, Xu JF, Zhang GJ. Clinical and genetic spectrum of primary ciliary dyskinesia in Chinese patients: a systematic review. Orphanet J Rare Dis. 2022;17(1):283.

43. Fliegauf M, Olbrich H, Horvath J, Wildhaber JH, Zariwala MA, Kennedy M, et al. Mislocalization of DNAH5 and DNAH9 in respiratory cells from patients with primary ciliary dyskinesia. Am J Respir Crit Care Med. 2005;171(12):1343–9.

44. Fassad MR, Shoemark A, Legendre M, Hirst RA, Koll F, le Borgne P, et al. Mutations in Outer Dynein Arm Heavy Chain DNAH9 Cause Motile Cilia Defects and Situs Inversus. Am J Hum Genet. 2018;103(6):984–94.

45. Zuccarello D, Ferlin A, Cazzadore C, Pepe A, Garolla A, Moretti A, et al. Mutations in dynein genes in patients affected by isolated non-syndromic asthenozoospermia. Hum Reprod. 2008;23(8):1957–62.

46. Wallmeier J, Nielsen KG, Kuehni CE, Lucas JS, Leigh MW, Zariwala MA, Omran H. Motile ciliopathies. Nat Rev Dis Primers. 2020;6(1):77.

47. Pan MM, Hockenberry MS, Kirby EW, Lipshultz LI. Male Infertility Diagnosis and Treatment in the Era of In Vitro Fertilization and Intracytoplasmic Sperm Injection. Med Clin North Am. 2018;102(2):337–47.

48. Esteves SC, Roque M, Bedoschi G, Haahr T, Humaidan P. Intracytoplasmic sperm injection for male infertility and consequences for offspring. Nat Rev Urol. 2018;15(9):535–62.

49. Long S, Fu L, Ma J, Yu H, Tang X, Han W, et al. O-253 Novel biallelic variants in DNAH1 cause multiple morphological abnormalities of sperm flagella with favorable outcomes of fertility after ICSI in Han Chinese males. Human Reproduction. 2023;38(Supplement_1).

50. Wambergue C, Zouari R, Fourati Ben Mustapha S, Martinez G, Devillard F, Hennebicq S, et al. Patients with multiple morphological abnormalities of the sperm flagella due to DNAH1 mutations have a good prognosis following intracytoplasmic sperm injection. Hum Reprod. 2016;31(6):1164–72.

51. Li Y, Sha Y, Wang X, Ding L, Liu W, Ji Z, et al. DNAH2 is a novel candidate gene associated with multiple morphological abnormalities of the sperm flagella. Clin Genet. 2019;95(5):590–600.

52. Gao Y, Tian S, Sha Y, Zha X, Cheng H, Wang A, et al. Novel bi-allelic variants in DNAH2 cause severe asthenoteratozoospermia with multiple morphological abnormalities of the flagella. Reprod Biomed Online. 2021;42(5):963–72.

53. Wei X, Sha Y, Wei Z, Zhu X, He F, Zhang X, et al. Bi-allelic mutations in DNAH7 cause asthenozoospermia by impairing the integrality of axoneme structure. Acta Biochim Biophys Sin (Shanghai). 2021;53(10):1300–9.

54. Zheng R, Sun Y, Jiang C, Chen D, Yang Y, Shen Y. A novel mutation in DNAH17 is present in a patient with multiple morphological abnormalities of the flagella. Reprod Biomed Online. 2021;43(3):532–41.

55. Huang F, Zeng J, Liu D, Zhang J, Liang B, Gao J, et al. A novel frameshift mutation in DNAH6 associated with male infertility and asthenoteratozoospermia. Front Endocrinol (Lausanne). 2023;14:1122004.

56. Li L, Sha YW, Xu X, Mei LB, Qiu PP, Ji ZY, et al. DNAH6 is a novel candidate gene associated with sperm head anomaly. Andrologia. 2018.

57. Liu Y, Niu M, Yao C, Hai Y, Yuan Q, Liu Y, et al. Fractionation of human spermatogenic cells using STA-PUT gravity sedimentation and their miRNA profiling. Sci Rep. 2015;5:8084.

58. Chang YF, Lee-Chang JS, Panneerdoss S, MacLean JA, 2nd, Rao MK. Isolation of Sertoli, Leydig, and spermatogenic cells from the mouse testis. Biotechniques. 2011;51(5):341–2, 4.

