## Supplementary figures and images for "DNAH3 deficiency causes flagellar inner dynein arm loss and male infertility in humans and mice"

### Figure S1

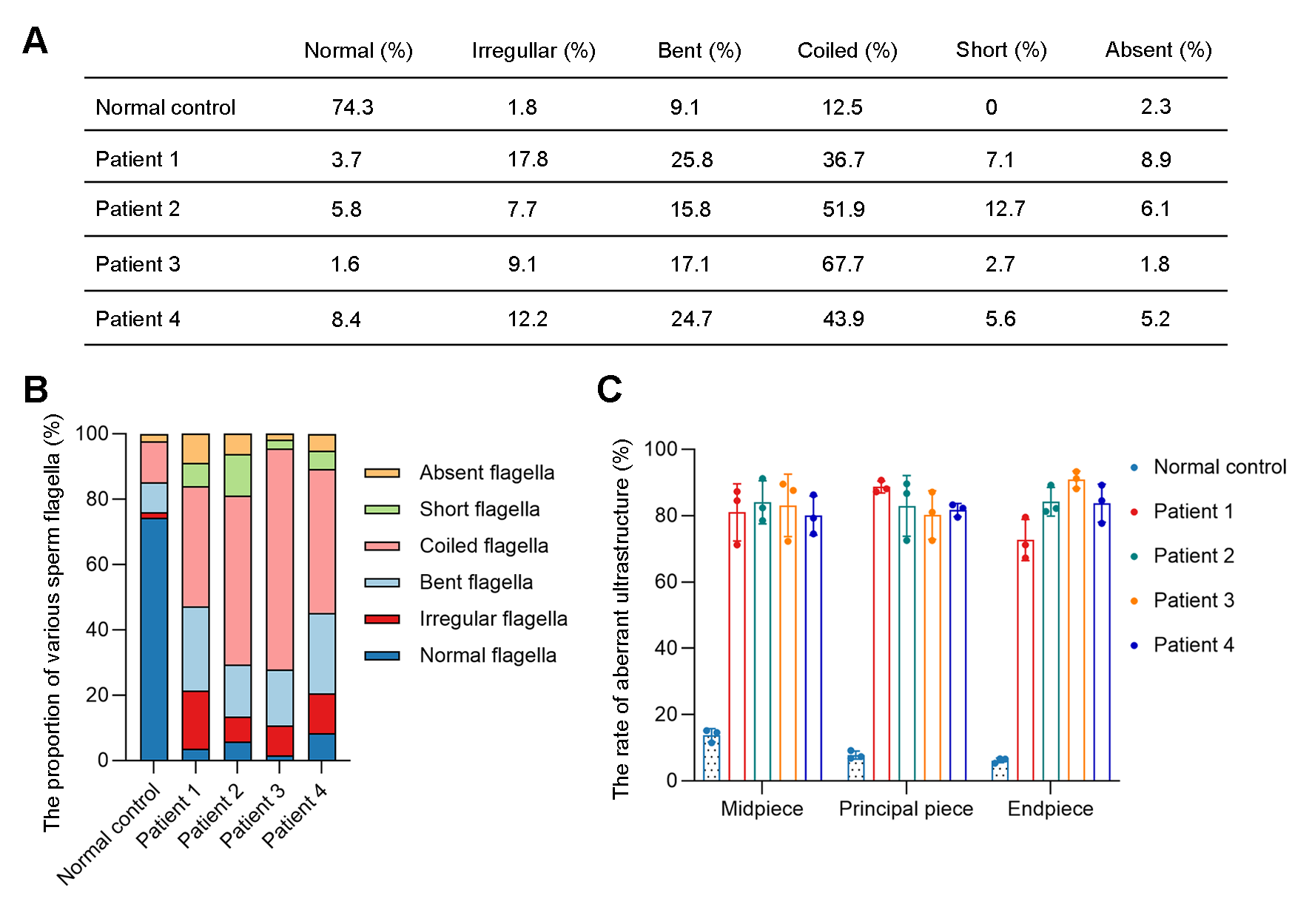

### Figure S2

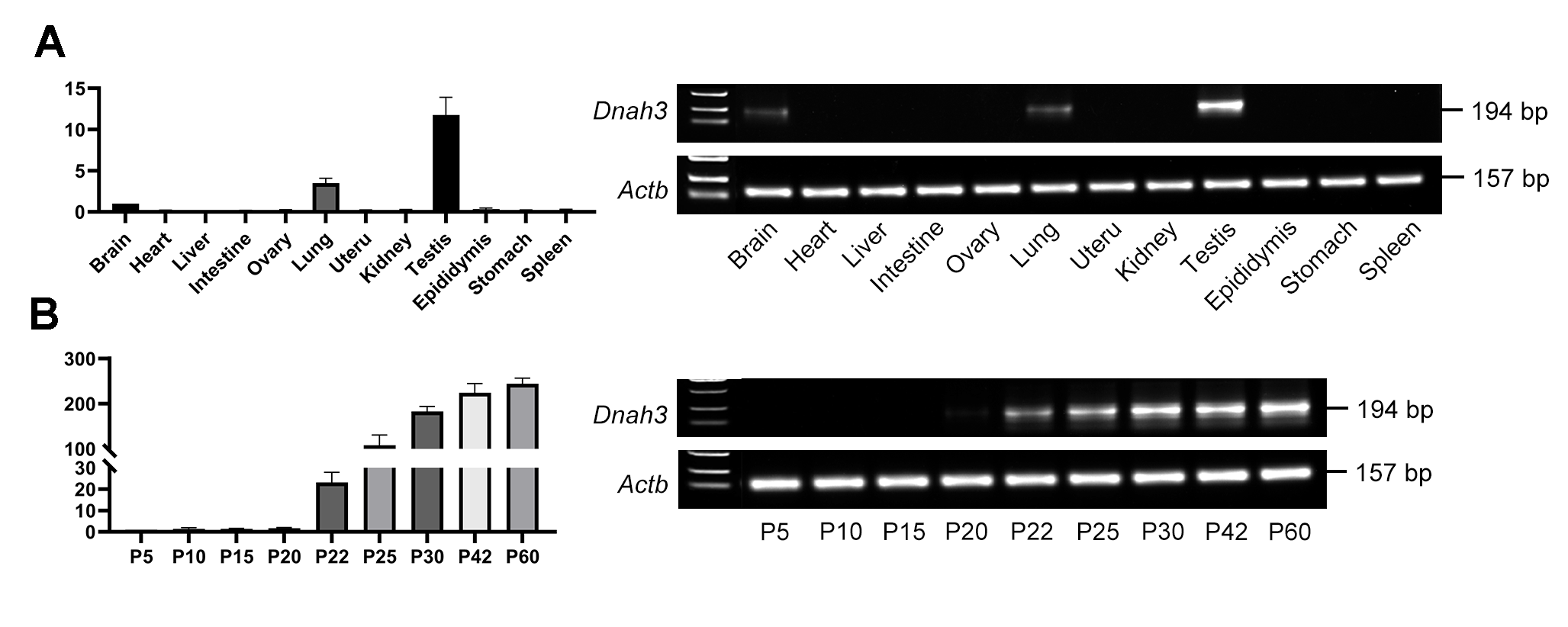

### Figure S3

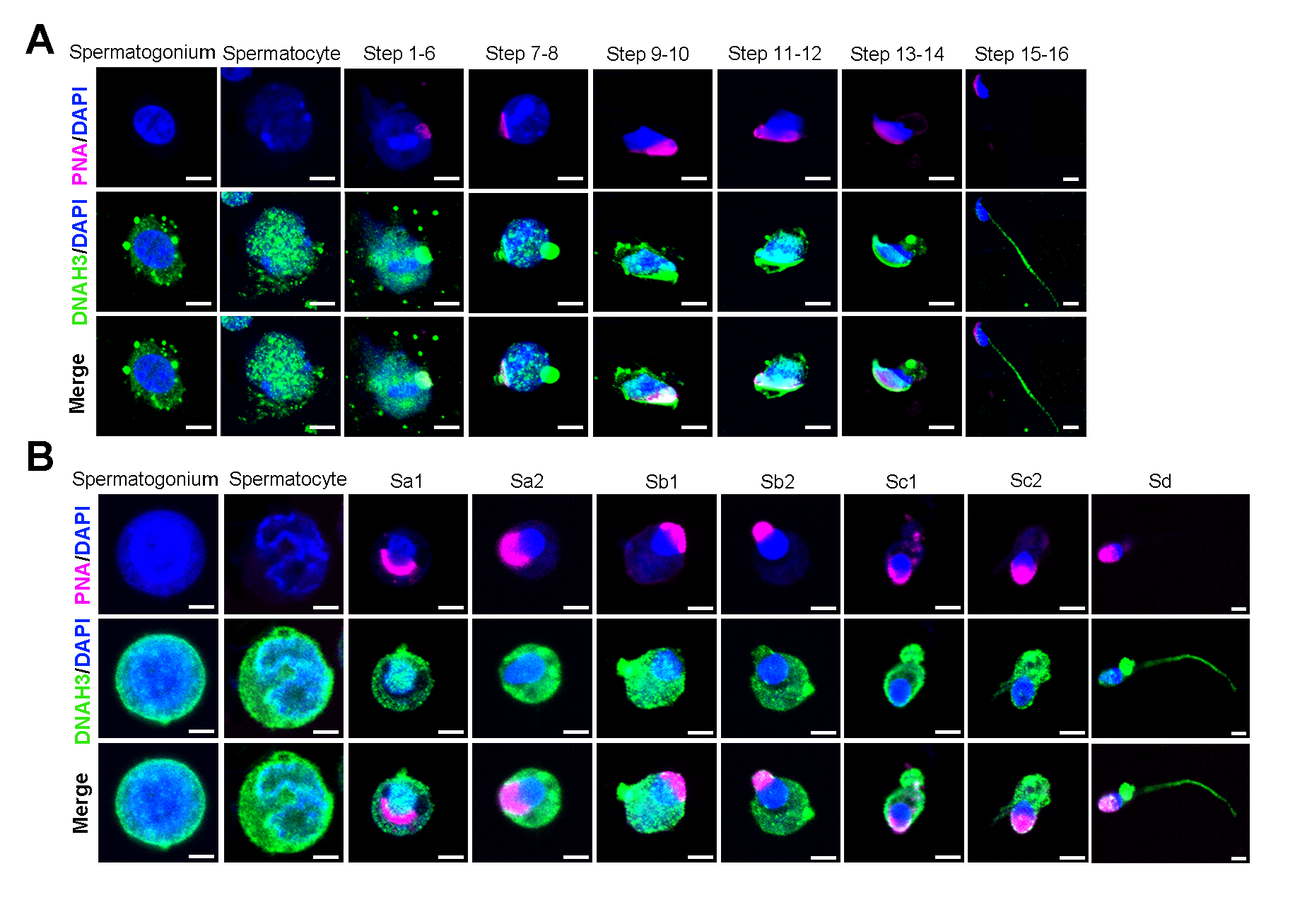

### Figure S4

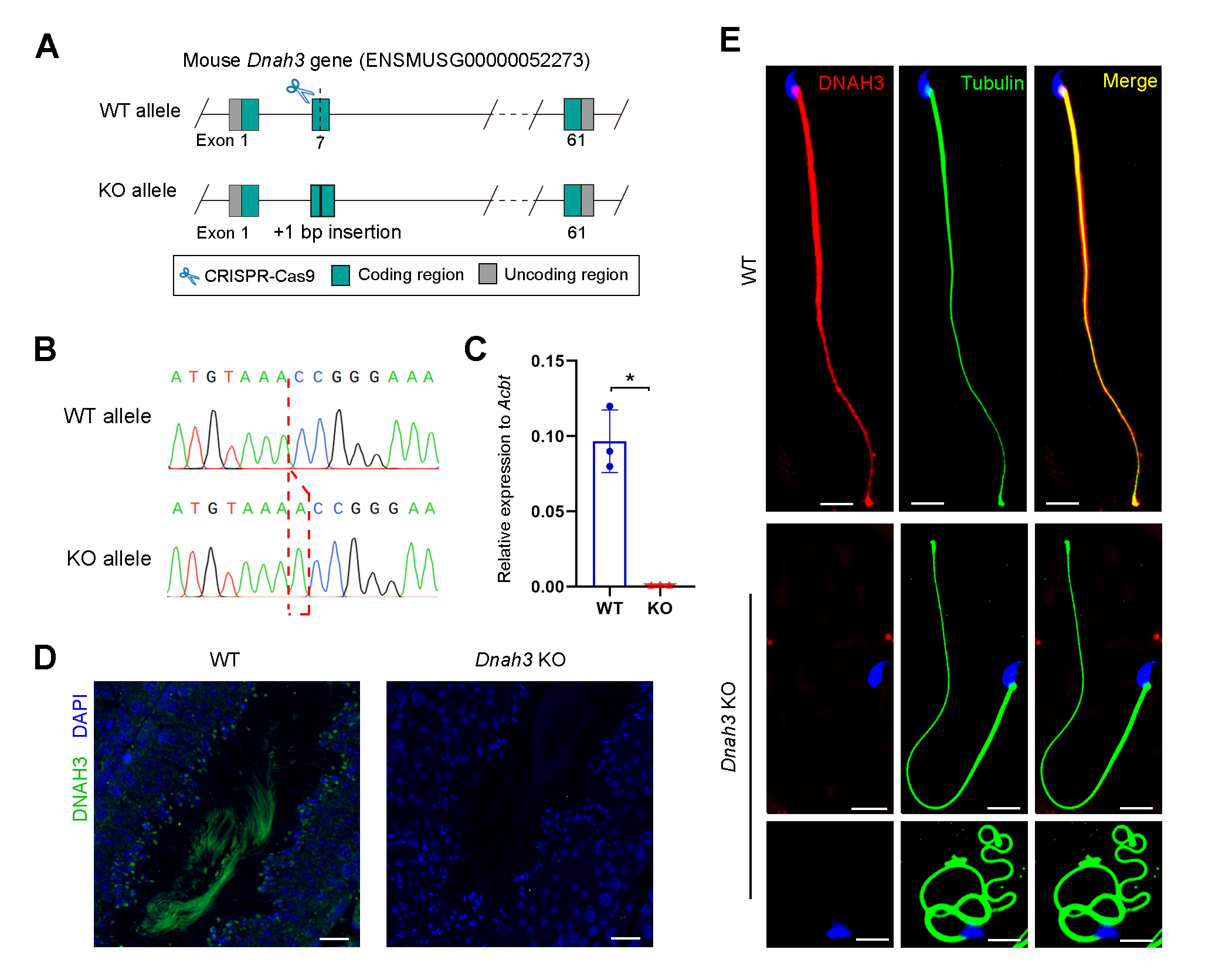

### Figure S5

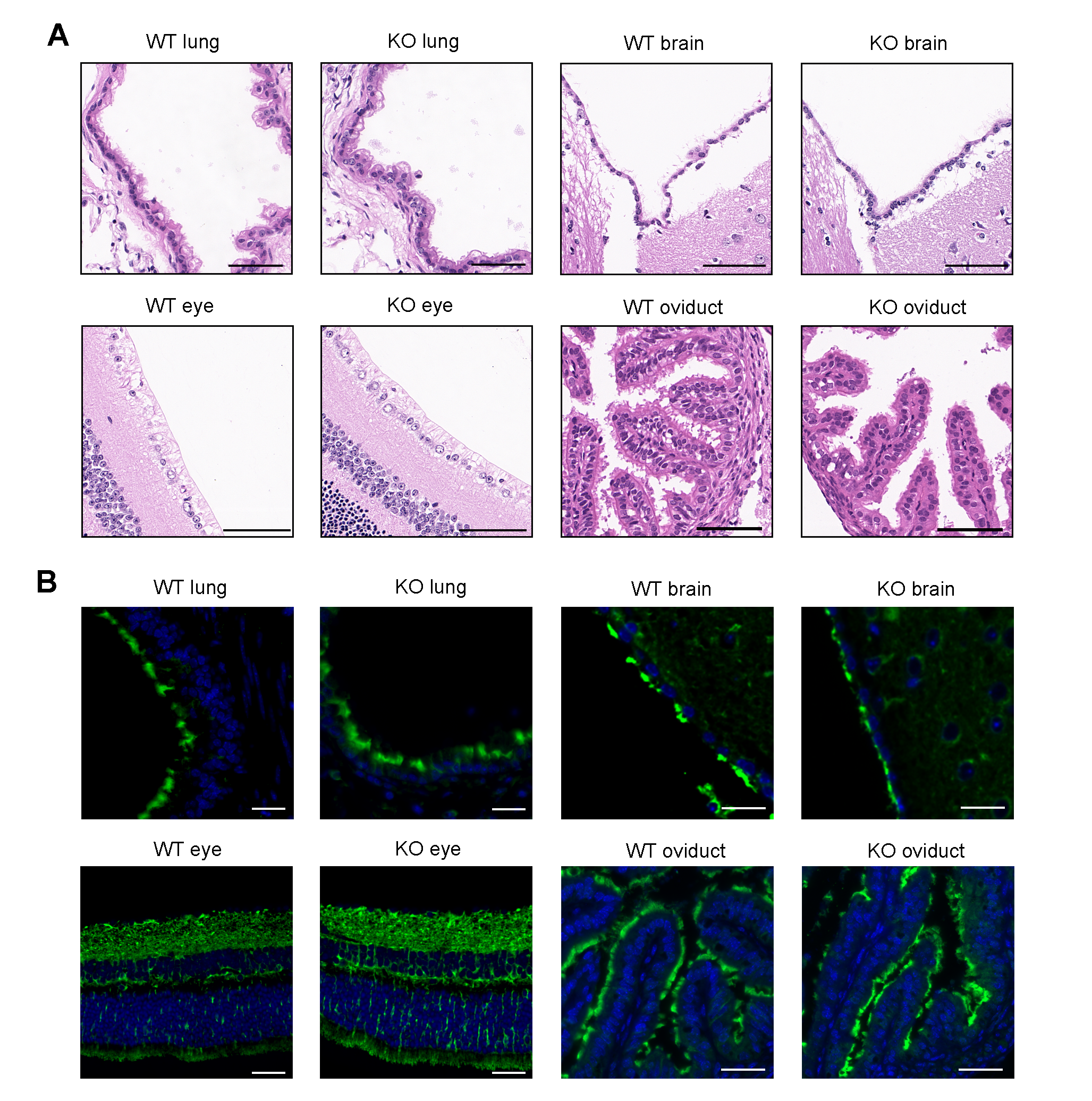

### Figure S6

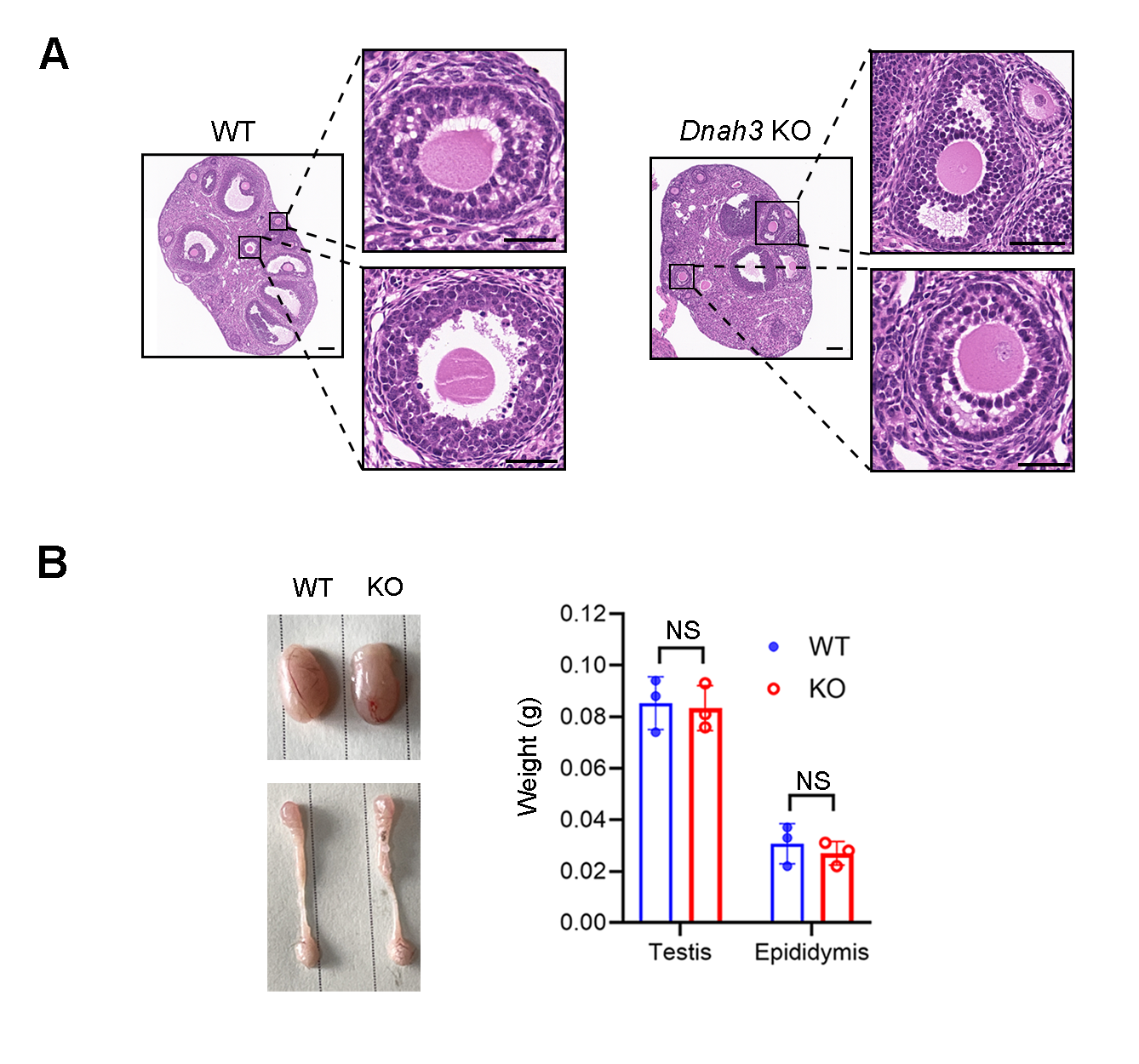

### Figure S7

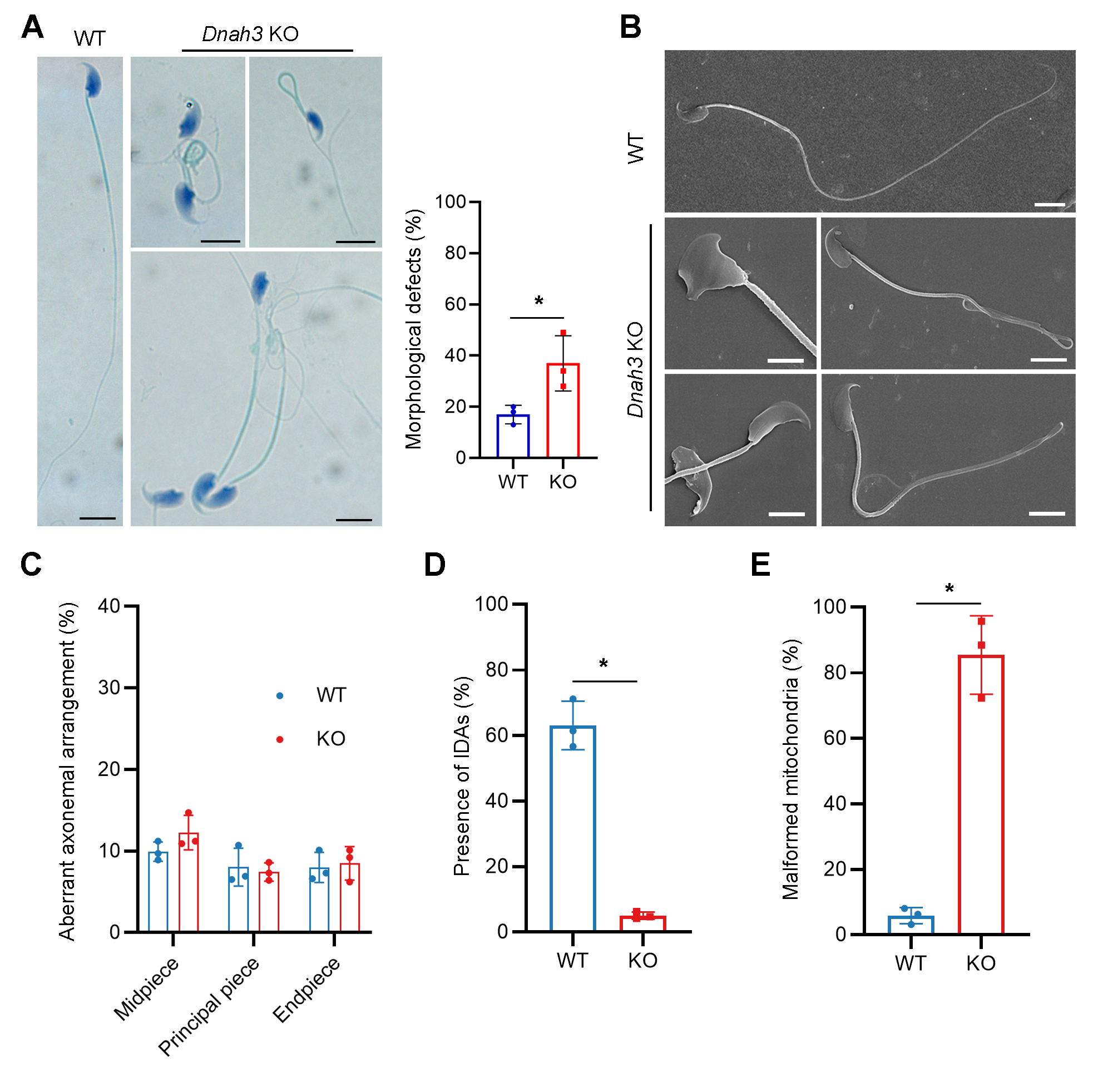

### Figure S8

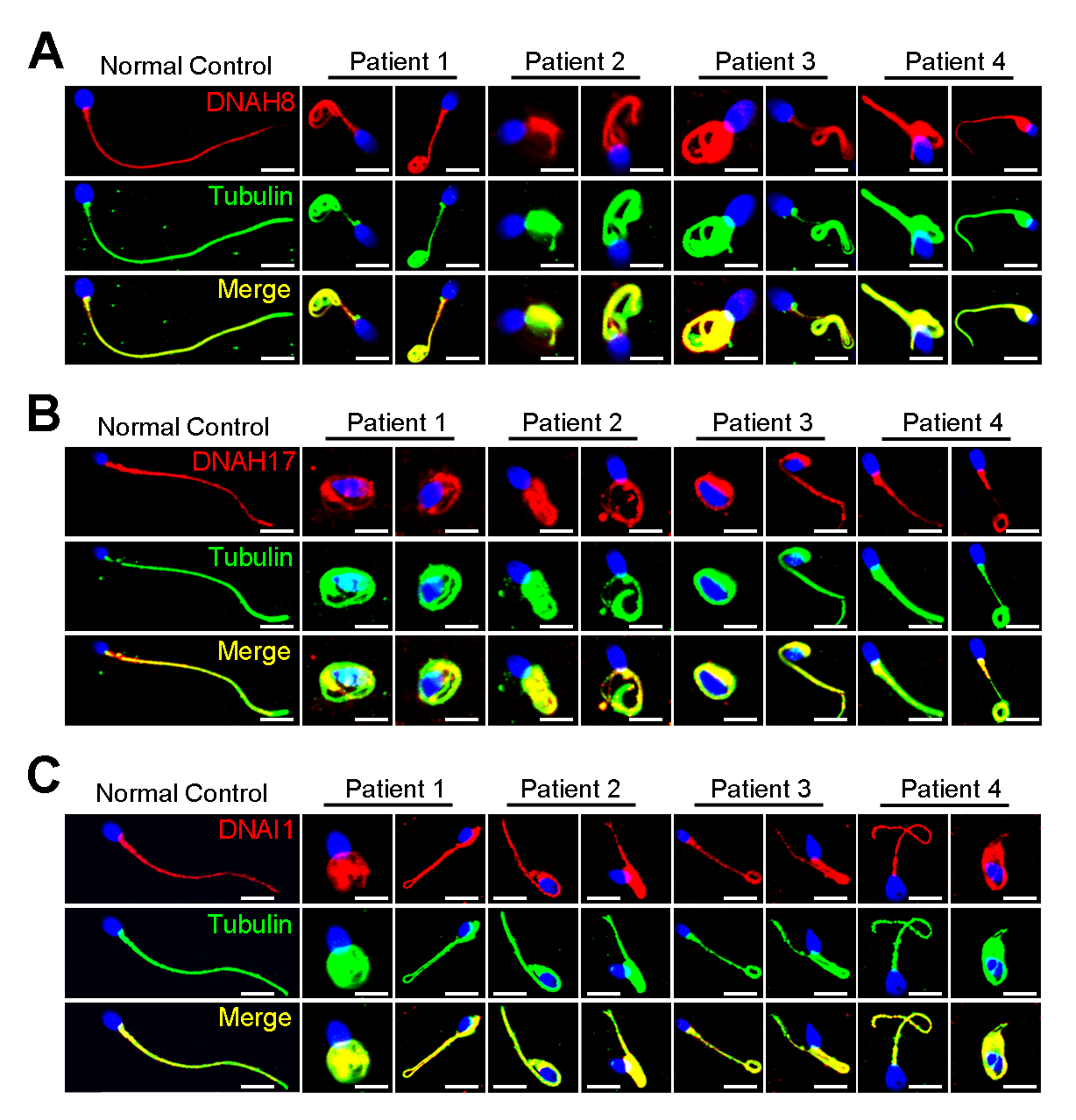

### Figure S9

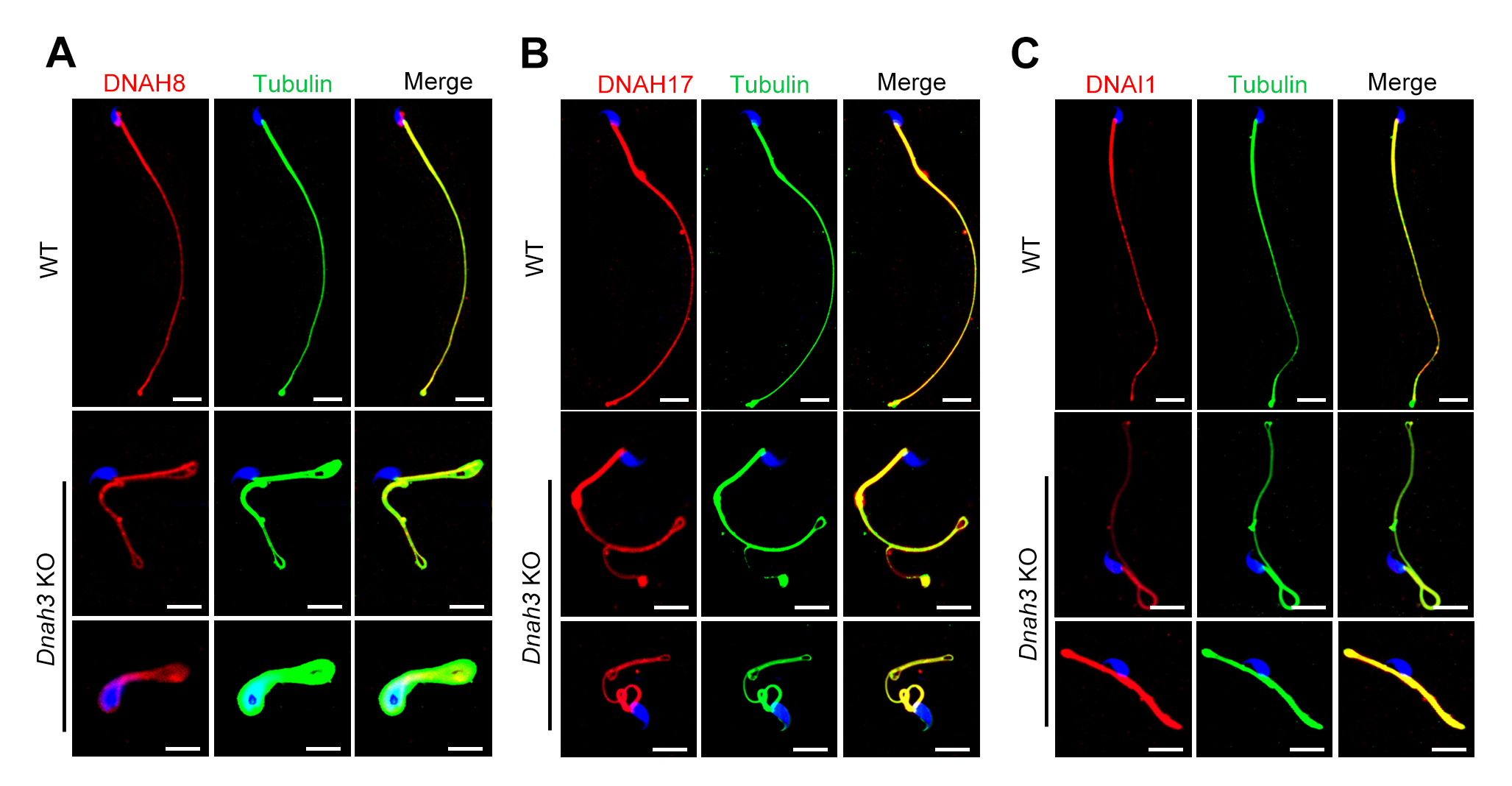
