## Supplementary material for "DNAH3 deficiency causes flagellar inner dynein arm loss and male infertility in humans and mice": Table S2

**Table S1**. **The primers used in the present study.**

| **Target** | **Forward primer (5’—3’)** | **Reverse primer (5’—3’)** | **Product (bp)** |
| --- | --- | --- | --- |
| **Sanger sequencing** | | | |
| c.3590C>G  c.3590C>T | TCTGTTATGGAGAAAAGAACCAA | GATGAAAAGTGATAAAAGAGAGTGG | 399 |
| c.4837G>T | TAACGCTGACCCCTGAATCTC | CAAGGTCCTGAACCGCTACAT | 876 |
| c.5587del | GCCCACCATATGGAGAAGAA | GAAATCAAGGGGCAGATGAA | 474 |
| c.10355C>T | TCCCTGGAGATCGAACGTAG | TGTGCCGTTCTCTGTTTGAG | 449 |
| c.2314C>T | CAGTGGGAAACCAAAGGAAA | CACACCACTCCTGTTGACCT | 392 |
| c.4045G>A | CAGAGTCTTCTTCCCCTGGA | TTGGAGAATGGGGGACCT | 368 |
| *Dnah3* | GAGAAGGGCATCAGTGAATT | TGTGGAGGTCCGTGGTTGAT | 624 |
| **qPCR** | | | |
| *Dnah3* | GGAGGTGATGATGCGAATTT | ATCGAGGGATGCTCTTGATG | 194 |
| *Actb* | CAGCTTCTTTGCAGCTCCTT | CACGATGGAGGGGAATACAG | 157 |
